# Context rescales a social action code in a hormone-sensitive network

**DOI:** 10.1101/2025.09.24.678271

**Authors:** Eartha Mae Guthman, Jorge M. Iravedra-Garcia, Lucy Sirrs, Alissa Le, Annegret L. Falkner

## Abstract

Deciding whether and when to engage in social interaction depends on external factors including the location of the interaction and the identity of the social partner (the social “context”) as well as internal factors such as an individual’s hormonal state. However, we lack a mechanistic understanding of how external and internal variables coordinate social action through networks of hormone-sensitive neurons. In particular, while gonadal hormones have long been suggested to coordinate territorial behaviors, direct evidence of how this coordination occurs has been lacking. To answer this, we combined large-scale neural recordings with large-scale unsupervised behavioral quantification^1^ to track neural activity longitudinally across a hormonal perturbation. We recorded neural population activity from neurons expressing the hormone receptor estrogen receptor alpha (ERα+) as well as from local ERα− neurons across the subcortical Social Behavior Network (SBN) and compared neural responses and behavior across social contexts with varied partners and territories. Using a comprehensive behavioral quantification strategy, we observe that patterns of social action and their underlying neural dynamics differentiate social partner and territory in both sexes. We find that each context has a unique behavioral action code, and that territory naturally rescales the partner-specific social action code in the hormonally intact state. However, when levels of circulating gonadal hormones are reduced, we observe that patterns of behavior during interactions in the home territory in males are disrupted, and that these changes can be rescued by testosterone replacement. Critically, hormonal perturbation disrupts territorial rescaling in a population-specific manner. Together, these data demonstrate how a loss of circulating hormones alters the relationship between social context and social action to disrupt context-specific social decision making.

## Introduction

The decision to engage in social behaviors depends on the social partner one encounters (the “who” of a social interaction) and the location of that encounter (the “where”). For example, male mice are more likely to initiate attacks towards male partners in their home territory than in the partner mouse’s territory but do not initiate attack towards female partners regardless of territory^2^. This combination of partner and location can be termed the social “context” for an interaction. Therefore, neural activity that determines the likelihood or timing of social behavior initiation must be sensitive to the conjunction of partner and location. In addition, social decision-making is also modulated by an individual’s internal state, including one’s hormonal state. However, it is unclear how social action codes are updated by context and whether the social context-specific patterning of behavior and its underlying neural codes are sensitive to hormonal state.

Activity in the brain’s subcortical social behavior network (SBN), in particular activity from hormone-sensitive neurons expressing estrogen receptor alpha (ERα), encodes critical information about partner type and identity^3–5^. The SBN is a hyper-connected set of subcortical regions largely in the hypothalamus, amygdala, and midbrain that are broadly conserved both anatomically and molecularly, in particular for receptors for gonadal hormones (e.g. estrogens and androgens)^6–10^. Numerous studies have established ERα+ neurons in the SBN as key mediators of territorial aggression, mating, and parenting behaviors^2,3,11–19^. Recordings from SBN ERα+ populations in males during social interactions in a mouse’s home territory have revealed many SBN populations are suppressed during female-directed behavior^4^, enabling the network to broadly distinguish male- and female-directed behaviors^3–5,14,18,20,21^. However, beyond the ability to distinguish the sex of a social partner, it has been unclear whether this network encodes additional social context variables, and whether social behavior encoding is specific to SBN ERα+ neural populations.

Modulation of a hormone-sensitive hypothalamic population within the social behavior network by estrogens has previously been shown to act as a gain control mechanism on neural activity to promote social approach behavior^22^, demonstrating that circulating gonadal hormones may play a generalized and flexible role in amplifying neural responses during proactive, or self-initiated, social action. Decades of research have demonstrated the necessity of testosterone (T) and testosterone-derived estrogen acting at neural androgen and estrogen receptors in territory establishment and defense across species^23–29^. However, a generalized model that directly links hormonal modulation to context-dependent social decision-making is lacking. For example, it is unclear whether gonadal hormones modulate specific subsets of critical behaviors, or suites of context-dependent behaviors.

To understand how social context and gonadal hormones influence social decision-making and its underlying neural coding across the social behavior network, we perform simultaneous, two-color imaging across 11 ERα+ and 11 ERα− SBN populations during social interaction^1^. We leverage recent advances in multi-animal pose-tracking^30,31^ to develop a comprehensive unsupervised analysis framework to identify and quantify patterns of mouse social behavior longitudinally across social contexts and hormonal state change. We demonstrate that circulating gonadal hormones are required for social context-dependent patterns of behavior in males but not females, and that T replacement rescues this context-dependent behavioral action patterning. Additionally, we show that territory serves as a natural amplification mechanism for behavior-associated neural activity across SBN populations, linearly rescaling the behavioral action code. Gonadectomy, which reduces levels of circulating gonadal hormones, disrupts this rescaling in a population specific manner. Together, our findings demonstrate how circulating T coordinates the specificity of a social action code to enable context-specific behavioral choices.

## Results

### Context influences fine-grain and coarse-grain patterns of social behavior

To map how social context influences patterns of social behavior, we took a comprehensive approach to map behavior across a variety of social contexts (Fig. 1A). Here we define context as the conjunction of the social partner the mouse is interacting with and the territory in which those interactions occur. Daily, each subject mouse (both males and females) interacted with 3 novel conspecific partners: An aggressive male CD-1 mouse (AGG+), a nonaggressive male BALB/c mouse (AGG-), and a female CD-1 mouse (FEM). Partner interactions could occur in each of two territories, the homecage of the subject mouse and the homecage of the partner mouse for a total of 6 social contexts each day. Context types were pseudorandomly presented across multiple days of behavior ensuring multiple trials for each context.

**Figure 1.**
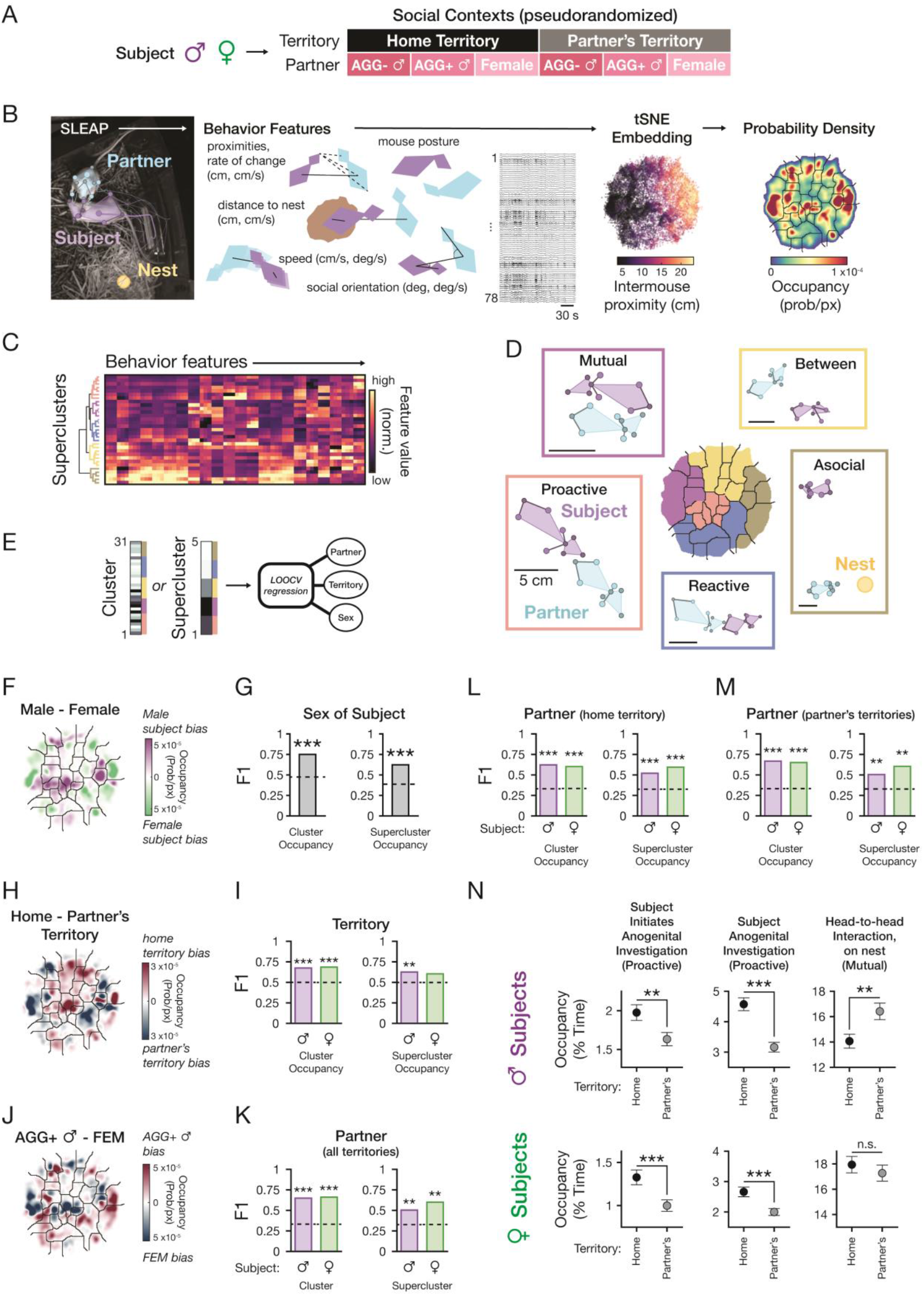
Patterns of behavior differentiate social context. A. Experimental timeline. Male or female subjects are presented with 6 different social contexts that vary partner and territory. B. Schematic demonstrating workflow from tracked pose to tSNE embeddings and derived probability density “Behavior Maps” with overlaid cluster boundaries. In the tSNE embedding, 0.1% of all frames plotted for visual clarity; N = 20.8 M frames, 2088 sessions, 58 mice (31 males, 27 females). C. Generation of superclusters via hierarchical clustering (Ward’s Method) of tSNE clusters using mean behavior feature values. D. Representative mouse postures during different Superclusters. Nodes shown: Nose, left and right ear, neck, left and right fore- and hindpaws, tailbase. Scale bar = 5 cm. E. Schematic of decoder workflow. F. Difference in occupancy between mean male and female subject behavior maps. G. Results of cluster and supercluster-based LOOCV subject sex decoder compared to shuffled session labels (*p*_cluster_ = 0, *p*_supercluster_ = 0, N = 558 male subject sessions, 31 mice, 486 female subject sessions, 27 mice). Dashed line shows mean of the shuffled label model. H. Difference between mean male subject behavior maps for home territory and partner’s territory interactions. I. Results of cluster and supercluster-based LOOCV territory decoder compared to shuffle (*p*_cluster_ = 0, *p*_supercluster_ = 0.008, male subject decoder, N = 279 male subject sessions per territory, 31 mice; (*p*_cluster_ = 0, *p*_supercluster_ = 0.054, female subject decoder, 243 female subject sessions per territory, 27 mice). J. Difference between mean male subject behavior maps for interactions with AGG+ male and FEM partner. K. Results of cluster and supercluster-based LOOCV partner decoder (all territories) compared to shuffle (*p*_cluster_ = 0, *p*_supercluster_ = 0, male subject decoder, N = 186 male subject sessions per partner; (*p*_cluster_ = 0, *p*_supercluster_ = 0.002, female subject decoder, 162 female subject sessions per territory). L. Same as K, but only using data from the subject’s home territory (*p*_cluster_ = 0, *p*_supercluster_ = 0, male subject decoder, N = 93 male subject sessions per partner, 31 mice; (*p*_cluster_ = 0, *p*_supercluster_ = 0, female subject decoder, 81 female subject sessions per territory, 27 mice). M. Same as K, but only using data from the partner’s territory (*p*_cluster_ = 0, *p*_supercluster_ = 0.006, male subject decoder, N = 93 male subject sessions per partner, 31 mice; (*p*_cluster_ = 0, *p*_supercluster_ = 0.002, female subject decoder, 81 female subject sessions per territory, 27 mice). N. Occupancy for example proactive and mutual behaviors. (Subject initiates anogenital investigation: *p*_MaleSubjects_ = 0.002; *p*_FemaleSubjects_ = 4.45 × 10^−4^. Subject anogenital investigation: *p*_MaleSubjects_ = 3.55 × 10^−9^; *p*_FemaleSubjects_ = 1.68 × 10^−4^. Head-to-head interaction: *p*_MaleSubjects_ = 0.004; *p*_FemaleSubjects_ = 0.44. All linear mixed effects models: N = 279 male subject sessions per territory, 31 mice; N = 243 male subject sessions per territory, 27 mice. Data presented as mean ± S.E.M.)

To identify common behaviors across contexts and track differences in those behaviors, we built on recent methods for unsupervised multi-animal behavioral classification^1,32–35^. We tracked the pose of the subject and partner mouse and the location of the nest using SLEAP^31^ (Fig. 1B, left). We converted raw pose to a set of 78 behavioral features, including relative social features (*e*.*g*., proximity and relative head angle), nonsocial features (*e*.*g*., distance from the nest), and egocentric variables (*e*.*g*., dimensionality reduced postural variables; see Table S1 for full list). We generated a nonlinear low dimensional embedding of these behavioral features using all data, recorded across subject sex and context (20.8 M frames), to create a two-dimensional behavior map. We segmented this behavior map using a watershed algorithm on the probability density of the behavior frames (Fig. 1B, right, see Methods). This approach results in a set of behavioral clusters that can be described as distinct action patterns and represent unique distributions of behavior features that co-occur (Fig. S1). We further grouped these fine-grain clusters into 5 coarse-grain “superclusters” using hierarchical clustering on a cluster by feature matrix of mean behavior feature values (Fig. 1C). These superclusters map to interpretable coarse-grain behavior classes (Fig. 1D; Movies S1-10) including proactive behavior (pink, social approach and interaction initiated or dominated by the subject mouse), mutual behavior (purple, social interactions between both mice without a clear initiator or leader), reactive behaviors (blue, social approach and interaction initiated or dominated by the partner mouse), asocial behaviors (brown, behavior lacking social interaction), and between behaviors (yellow, behavioral actions while the mice transition between social and asocial behavior or vice-versa). These superclusters are broadly consistent with categories of social behavior observed using a similar unsupervised technique to quantify rat social behavior^35^.

Next, we tested whether patterns of fine-grain (clusters) or coarse-grain (superclusters) behavioral action predict social context. To do this, we generated occupancy vectors of percent time spent in each cluster on a session-by-session basis and tested whether we can predict subject sex or context variables (partner, territory) using leave-one-out logistic regression to train a decoder (Fig. 1E). We find that we can accurately decode subject sex (Fig. 1G) using both fine- and coarse grain-methods. By comparing the behavior maps and behavior occupancy we observe a clear bias in males for proactive, or self-initiated, social behaviors (Fig. 1F; Fig S2A).

In addition, we find that we can significantly decode territory (Fig. 1H-I, purple) and partner type (Fig. 1J-K, purple) using both fine- and coarse-grain approaches in males. In females, territory and partner can be decoded from cluster occupancy, but only partner can be decoded from superclusters above chance level. Further, we find we can decode partner type individually in each territory for males and females (Fig. 1L-M). This approach demonstrates that coarse-graining captures important behavior differences and that behavior differs at several levels of the behavior hierarchy.

By comparing the behavior maps and behavior occupancy across contexts, we observe that proactive behaviors are most prevalent in the home territory and during FEM interactions in both males and females (Fig. 1H, J, N; Fig. S2B-C). In contrast to proactive behaviors, we find that mutual behaviors are more prevalent in the partner’s territory in males but not females and more prevalent during AGG+ and FEM interactions in both sexes (Fig. 1H, N; Fig. S2D-E). In females, we find that the behavioral landscape does not vary significantly with estrous stage (Fig. S3). Overall, these data suggest that patterns of behavior are highly distinguishable across contexts in both sexes.

### Subcortical hormone-sensitive networks encode partner, territory and social action in a distributed code

To test whether patterns of social behavior are encoded in a context-specific manner, we recorded and compared brain-wide neural activity across all behaviors across all 6 contexts. We targeted the conserved subcortical SBN, which richly expresses the hormone receptor ERα across a network of highly connected regions in the forebrain, hypothalamus, amygdala, and midbrain (Fig. 2A). Activation of ERα expressing neurons in a variety of sites in this network are well known to promote specific forms of proactive behaviors^2,3,11–17,19^. To record from neurons expressing ERα (ERα+) and neurons not-expressing ERα (ERα−) in the same region, we used a dual viral strategy in Esr1-Cre male and female subject mice: We stereotactically injected both a Cre-dependent red calcium indicator^36^ (jRCaMP1b) and a Cre-excluding^37^ green calcium indicator (GCaMP6f) at each of 11 sites in the SBN (Fig. S4). These calcium indicators are known to have relatively matched kinetics^36^. We next implanted a custom array of 200 μm fibers over these sites and recorded neural activity using alternating pulsed 560nm and 470nm LEDs interleaved with an isosbestic wavelength (Fig. 2B). We validated that opposing LEDs activate very little calcium indicator (Fig. 2C) and performed motion correction on this signal using two-color motion artifact correction^38^ (Fig. 2D). Next, we recorded activity simultaneously across all ERα+ and ERα− populations during social behavior and assign timepoints with cluster and supercluster behavior labels (Fig. 2E-F) in both male and female subject mice. To compare neural activity across contexts, we projected the neural activity from each signal onto our master behavior embedding (Fig. 2G).

**Figure 2.**
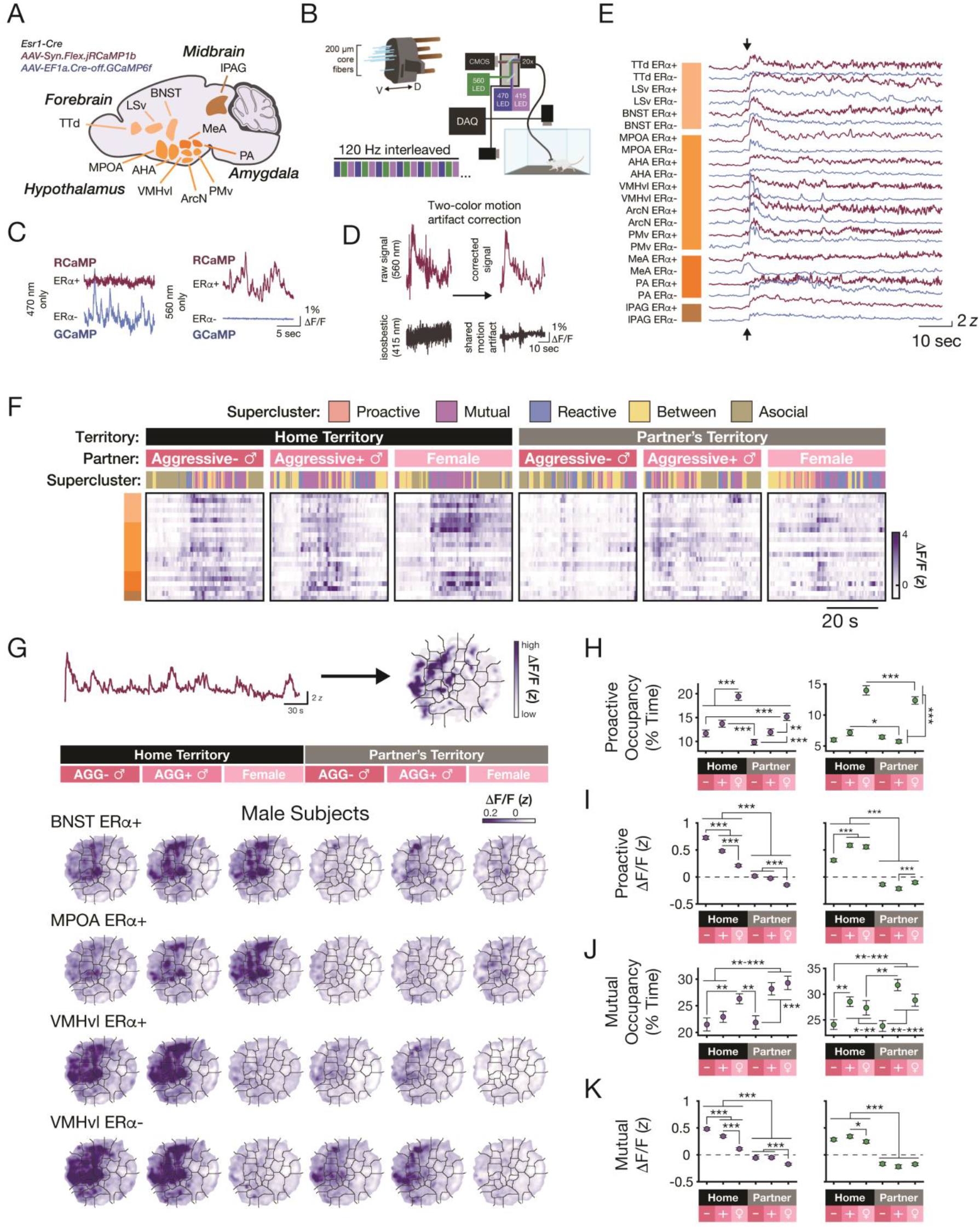
Context differentiates patterns of neural activity in the social behavior network. A. Viral strategy schematic and targeted brain regions. Abbreviations, TTd: Tenia Tecta, dorsal part; LSv: Lateral Septum, ventral part; BNST: Bed Nucleus of the Stria Terminalis; MPOA: Medial Preoptic Area; AHA: Anterior Hypothalamic Area; VMHvl: Ventromedial Hypothalamus, ventrolateral part; ArcN: Arcuate Nucleus of the Hypothalamus; PMv: Ventral Premammillary Hypothalamus; MeA: Medial Amygdala; PA: Posterior Amygdala; lPAG: Lateral Periaqueductal Gray. B. Multisite photometry schematic. C. Independence of RCaMP and GCaMP signals. D. Example detailing use of Two-color motion artifact correction strategy used. E. Example SBN-wide recording at time of partner entrance (arrowhead). Color bars correspond to brain regions (forebrain, hypothalamus, amygdala, midbrain) of recorded population. Red traces are RCaMP (ERα^+^) signals, and blue traces are GCaMP (ERα^−^) signals. F. 45s of SBN-wide neural activity from each of the 6 sessions on a single experimental day from a representative male mouse. G. Top: Schematic detailing the process by which neural signals are projected onto the behavior maps. Bottom: Mean heatmaps for example SBN populations (N = 36 activity maps per context, 12 intact male mice). H. Proactive supercluster occupancy across social contexts. Left: Male subjects (Linear Mixed Effects Model, α_FDR_ = 0.0271, N = 93 sessions, 31 mice). Right: Female subjects (Linear Mixed Effects Model, α_FDR_ = 0.0312, N = 81 sessions, 27 mice). I. Mean SBN neural activity during proactive behaviors across social contexts. Left: Male subjects (Linear Mixed Effects Model, α_FDR_ = 0.0479, N = 770-777 populations/context, 12 mice). Right: Female subjects (Linear Mixed Effects Model, α_FDR_ = 0.0438, N = 322-325 populations/context populations, 5 mice). J. Mutual supercluster occupancy across social contexts. Left: Male subjects (Linear Mixed Effects Model, α_FDR_ = 0.0250, N = 93 sessions, 31 mice). Right: Female subjects (Linear Mixed Effects Model, α_FDR_ = 0.0271, N = 81 sessions, 27 mice). K. Mean SBN neural activity during mutual behaviors across social contexts. Left: Male subjects (Linear Mixed Effects Model, α_FDR_ = 0.0479, N = 770-777 populations/context, 12 mice). Right: Female subjects (Linear Mixed Effects Model, α_FDR_ = 0.0354, N = 322-325 populations/context, 5 mice). Data presented as mean ± S.E.M. Linear Mixed Effects Model formulas can be found in Methods. *: *p* < α_FDR_, **: *p* < 0.01, ***: *p* < 0.001.

Using this method, we observe that neural activity in both ERα+ and ERα− populations exhibits activity gradients that vary across partner, territory, and behavior. We observe systematic but complex variation in the relationship between neural activity and occupancy for social behavior clusters across contexts, specifically for proactive and mutual behaviors (Fig. 2H-K), suggesting that each context contains a distinct mapping of neural activity to behavior. However, high levels of activity do not always predict high levels of that behavior. For example, in males, we observe that within a territory type, there is no consistent relationship between neural activity and occupancy, and, in some cases, the relationship is inverted, while in females this relationship is even more complex (Fig. 2H-K, Table S2). This demonstrates that SBN neural population activity has context-specific patterns of activity that do not simply predict more or less of a given behavior.

To test more directly whether patterns of activity differentiate social contexts, irrespective of behavior, we trained a series of 10-fold cross validated (CV) linear decoders to predict territory, partner, or behavior on a moment-to-moment basis (Fig. 3A). This is akin to asking whether the instantaneous activity across the SBN signals information about these three variables simultaneously, independent of the other variables. For both male and female subjects, we find we can decode which territory interactions occurred in and which partner the subject interacted with significantly better than a control with shuffled labels (Fig. 3B). To test the contributions of ERα+ or ERα− populations to each of these decoders, we iteratively shuffled either ERα+ or ERα− populations, and computed the F1 loss, a measure of how much the decoder accuracy suffers. We find no significant difference in F1 loss for any decoder between these populations, indicating that they both contribute to SBN encoding of social context (Fig. 3B). To quantify the contribution of each neural population to the decoder we retrained the decoder after cumulatively shuffling SBN population activity. We rank-ordered and shuffled populations in order of descending decoder weight, starting with the signal with the largest weight (Fig. 3C-D; Fig. S5). If SBN encoding of context specific variables is modular (e.g. a handful of brain regions encode territory, while many do not participate), we would expect a drop in F1 to chance levels before all populations are shuffled. In contrast, if SBN encoding of these variables is distributed, we would expect a more gradual decrease in decoding accuracy as more populations are removed. We find that for all context related variables, encoding is widely distributed across the SBN. However, we observe that a handful of SBN populations, including LSv ERα+ and VMHvl ERα+ and ERα− neurons consistently have the largest decoder weights, indicating that they may play outsized roles in SBN encoding of social context.

**Figure 3.**
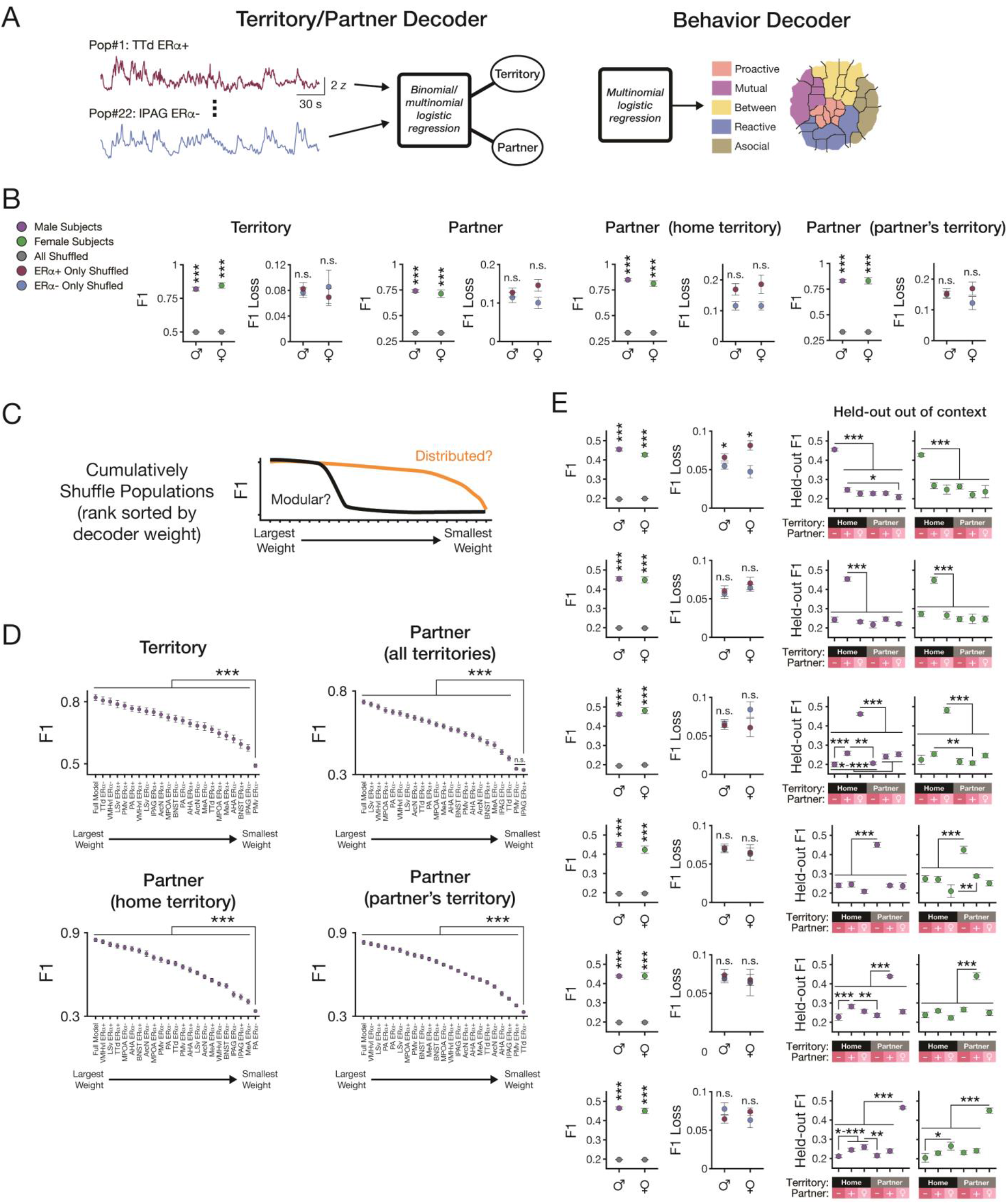
Social Behavior Network encodes social context and behavior in a distributed manner and at a moment-to-moment scale. A. Decoders for territory, partner, and behavior. B. Territory and partner decoder results showing 10-fold CV decoder F1 score (left) and F1 Loss following ERα+/− shuffle (right). Territory, base model: *p*_Male_ = 2.51 × 10^−10^, *p*_Female_ = 7.32 × 10^−5^ Territory, F1 Loss: *p*_Male_ = 0.72, *p*_Female_ = 0.60. Partner (all territories), base model: *p*_Male_ = 3.23 × 10^−12^, *p*_Female_ = 5.23 × 10^−4^. Partner (all territories), F1 Loss: *p*_Male_ = 0.53, *p*_Female_ = 0.14. Partner (home territory), base model: *p*_Male_ = 3.19 × 10^−13^, *p*_Female_ = 6.52 × 10^−5^. Partner (home territory), F1 Loss: *p*_Male_ = 0.10, *p*_Female_ = 0.14. Partner (partner’s territory), base model: *p*_Male_ = 2.31 × 10^−12^, *p*_Female_ = 9.13 × 10^−5^. Partner (partner’s territory), F1 Loss: *p*_Male_ = 0.91, *p*_Female_ = 0.28. C. Predicted outcomes of cumulatively shuffled 10-fold CV decoder. D. Cumulatively shuffled 10-fold CV territory and partner decoders in male mice. Linear mixed effects model used to compare decoding in the fully shuffled condition (i.e. all populations from largest through smallest weight shuffled) against all other runs of the cumulatively decoder. Territory decoding: PMv ERα− vs. all other pops, all *p* < 1.56 × 10^−10^. Partner decoding (all territories): lPAG ERα+ vs. PMv ERα+, *p* = 0.62; lPAG ERα+ vs. all other pops, all *p* < 5.00 × 10^−7^. Partner decoding (home territory): PA ERα− vs. all other pops, all *p* < 4.37 × 10^−7^. Partner decoding (home territory): PA ERα− vs. all other pops, all *p* < 2.81 × 10^−4^. E. 10-fold CV context-specific action decoder results. Each row, left to right: Base decoder F1 score (real vs. shuffle), F1 Loss following ERα+/− shuffle, F1 score for base decoder tested on held-out data from each of the 6 social contexts. Context order, top to bottom: Home territory, AGG-; Home territory, AGG+; Home territory, FEM; Partner’s territory, AGG-; Partner’s territory, AGG+; Partner’s territory, FEM. Base model F1 score real vs. shuffle, F1 Loss ERα+ vs. ERα− were compared using t-tests, and Held-out F1 score was compared using linear mixed effects models. All statistics in Table S3. N = 12 male models, 5 female models. All statistical tests are paired t-tests unless otherwise stated. Data presented as mean ± S.E.M. Linear Mixed Effects Model formulas can be found in Methods. *: *p* < 0.05, **: *p* < 0.01, ***: *p* < 0.001.

### The social behavior network encodes patterns of behavior using context-specific schema

Next, we tested whether each context has a unique action coding schema or uses a shared neural profile for social action (Fig. 3E). First, we trained an action decoder on SBN neural activity from each context to predict supercluster behavior on a moment-to-moment basis. We built models using our coarse-grain approach because our fine-grain approach resulted in more behavior clusters than recorded neural populations. Within each context, we find we can decode behavior significantly better than a shuffle control in both males and females (Fig. 3E, left). We find that both ERα+ and ERα− populations contribute similarly to action decoding, except for in the home territory-AGG-partner context, for which shuffling ERα+ populations results in increased F1 loss relative to ERα− populations in both males and females (Fig. 3E, middle). Similar to context encoding, we find that the SBN encodes behavioral action in a distributed manner, regardless of context (Fig. S5). Finally, when we test each within-context decoder on held-out out-of-context neural activity, we find a significant degradation in decoding accuracy to near chance levels (Fig. 3E, right). This demonstrates that each context has a unique profile of behavior-specific neural activity.

### Home territory rescales context-specific neural activity

As each context has a unique action code, we sought a simple model that would explain the change in neural activity across contexts. We find the relationship between mean neural population response for the same behavior performed in the home territory and the partner’s territory is well described by a linear model with a single coefficient and intercept (Fig. 4A-D). This model performs better than chance across and within partner types (Fig. 4D, Fig. S6A). We find that fit coefficients and intercepts are uniformly positive, with better fits and greater coefficients in males compared to females (Fig. 4D-F, Fig. S6B). This indicates that home territory rescales and shifts neural activity levels relative to the partner’s territory with greater rescaling occurring in males, as observed in context-specific tuning curves (Fig. 4B). In addition, we observe that residuals for individual behaviors are positive for proactive behaviors and negative for nonsocial behaviors, suggesting greater territorial rescaling of neural activity for these specific behaviors (Fig. 4E). Importantly, this rescaling is not due to changes in neural activity during specific behaviors such as aggression; rescaling of the behavior code, or the vector of mean neural population activity across behavior clusters, remains well described by a linear model fit without the inclusion of any proactive behaviors (Fig. S6C), indicating that this is a translation applied to all behavioral action. Finally, unsupervised clustering of the fit coefficients and intercepts reveals a subnetwork of brain regions that undergo greater rescaling during home interactions with males (including both ERα+ and ERα− VMHvl populations, ERα+ AHA, ERα− BNST, and ERα− PA populations; Fig. 4F). Notably, we observe this subnetwork independently in both males and females, suggesting a generalized sharing of information about male partners in this subnetwork. Together, these data indicate that being in one’s home territory acts as a natural amplification mechanism to rescale the neural responses to all behaviors, independent of partner type.

**Figure 4.**
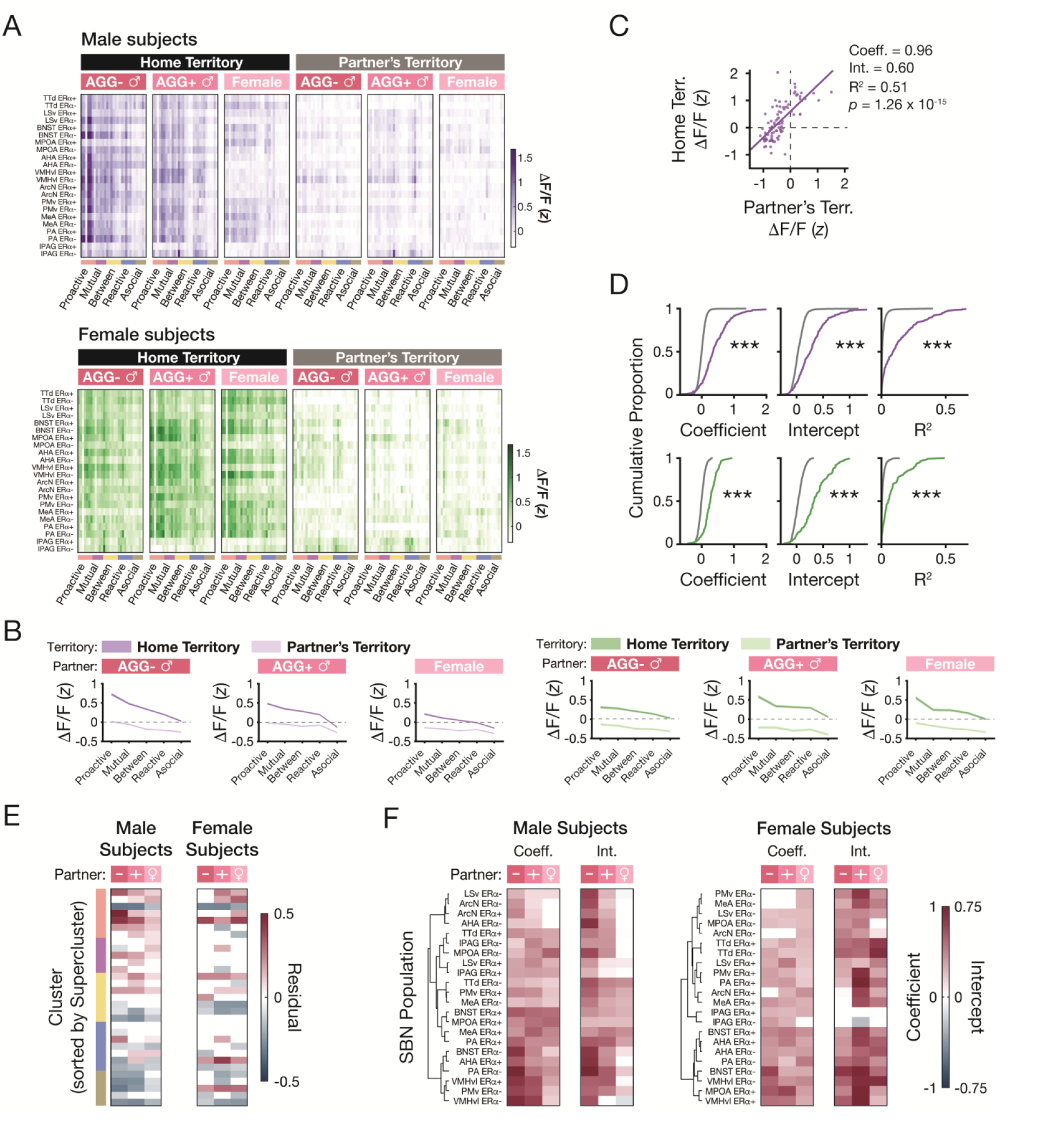
Home territory functions as a natural gain control mechanism. A. Heatmaps of mean neural activity across all recorded populations and behavior clusters in each social context. Top: Male subjects. Bottom: Female subjects. B. Mean supercluster tuning curves during home and partner’s territory interactions with AGG+, AGG-, and FEM partners. Left: Male subjects. Right: Female subjects. C. Example cross-territory regression from the MPOA ERα+ population of a representative male mouse. Each data point represents the z-scored neural activity in the home and partner’s territories for a given partner-behavior cluster pairing. D. Distributions of regression coefficients, intercepts, and R^2^ in male (top) and female (bottom) subjects compared to shuffled cluster labels. Male: *p*_Coefficient_ = 2.21 × 10^−88^, *p*_Intercept_ = 4.09 × 10^−54^, *p*_R-Squared_ = 5.79 × 10^−70^, N = 257 populations, 12 mice; Female: *p*_Coefficient_ = 1.08 × 10^−40^, *p*_Intercept_ = 5.01 × 10^−41^, *p*_R-Squared_ = 2.30 × 10^−29^, N = 107 populations, 5 mice; Kolmogorov-Smirnov tests. E. Heatmap of mean regression residuals across partner identities and behavior clusters. Only clusters with residuals significantly different from shuffle are shown. Linear Mixed Effects Model, *p* < α_FDR_; Male subject, AGG-partner α_FDR_: 0.0387, N = 7,790 population residuals, 31 clusters, 12 mice; Male subject, AGG+ partner α_FDR_: 0.0355, N = 7,790 population residuals, 31 clusters, 12 mice; Male subject, FEM partner α_FDR_: 0.0355, N = 7,760 population residuals, 31 clusters, 12 mice; Female subject, AGG-partner α_FDR_: 0.0194, N = 3,245 population residuals, 31 clusters, 5 mice; Female subject, AGG+ partner α_FDR_: 0.0274, N = 3,274 population residuals, 31 clusters, 5 mice; Female subject, FEM partner α_FDR_: 0.029 N = 3,286 population residuals, 31 clusters, 5 mice. F. Heatmap of mean regression coefficients and intercepts across partner identities and neural populations in male (left) and female (right) subjects. Neural populations are clustered based on the combined coefficient and intercept heatmaps. Only populations with coefficients or intercepts significantly different than shuffle are shown. Linear Mixed Effects Model, *p* < α_FDR_; Male subject, AGG-partner Beta α_FDR_: 0.05, Intercept α_FDR_: 0.05, N = 252 populations, 12 mice; Male subject, AGG+ partner Beta α_FDR_: 0.05, Intercept α_FDR_: 0.05, N = 252 populations, 12 mice; Male subject, FEM partner Beta α_FDR_: 0.0432, Intercept α_FDR_: 0.0409, N = 251 populations, 12 mice; Female subject, AGG-partner Beta α_FDR_: 0.0409, Intercept α_FDR_: 0.0432, N = 106 populations, 5 mice; Female subject, AGG+ partner Beta α_FDR_: 0.0432, Intercept α_FDR_: 0.05, N = 107 populations, 5 mice; Female subject, FEM partner Beta α_FDR_: 0.0477, Intercept α_FDR_: 0.05, N = 106 populations, 5 mice. Data presented as mean ± S.E.M. Linear Mixed Effects Model formulas can be found in Methods. *: *p* < α_FDR_, **: *p* < 0.01, ***: *p* < 0.001.

### Context-specific patterns of social behavior in males are disrupted by gonadectomy and rescued by testosterone replacement

Prior work demonstrated gonadal hormones can adjust the neural response gain in a subset of ERα+ MPOA neurons^22^. We hypothesized the territorial rescaling we observe across the SBN might also require hormonal modulation. A reduction in circulating gonadal hormones reduces specific social behaviors, such as aggression^39^. However, whether these effects are limited to specific social contexts or actions remains unknown. Here, we used gonadectomy (GDX) in adulthood, also known as orchiectomy in males or ovariectomy in females, to directly perturb circulating gonadal hormones and tested how this perturbation affected patterns of behavioral action across our 6 partner-territory contexts (Fig. 5A). We compared behavior in animals before and after GDX or sham GDX. Given the complexity of the endogenous hormonal milieu regardless of sex^40,41^, we tested whether merely replacing T (6.6 μg/hr) following GDX is sufficient to recover pre-GDX patterns of behaviors. We focused on T because conspecific aggression requires T in both male and female rodents^25,42^. To provide minimally invasive T replacement, we used osmotic minipumps to deliver T rather than daily injections. We compared the behavior profiles across 3 groups (Fig. 5A): The sham control group, which receives sham surgery and no hormone replacement (oil), the GDX group, which receives GDX and no hormone replacement (oil), and the T group, which receives both GDX and T replacement.

**Figure 5.**
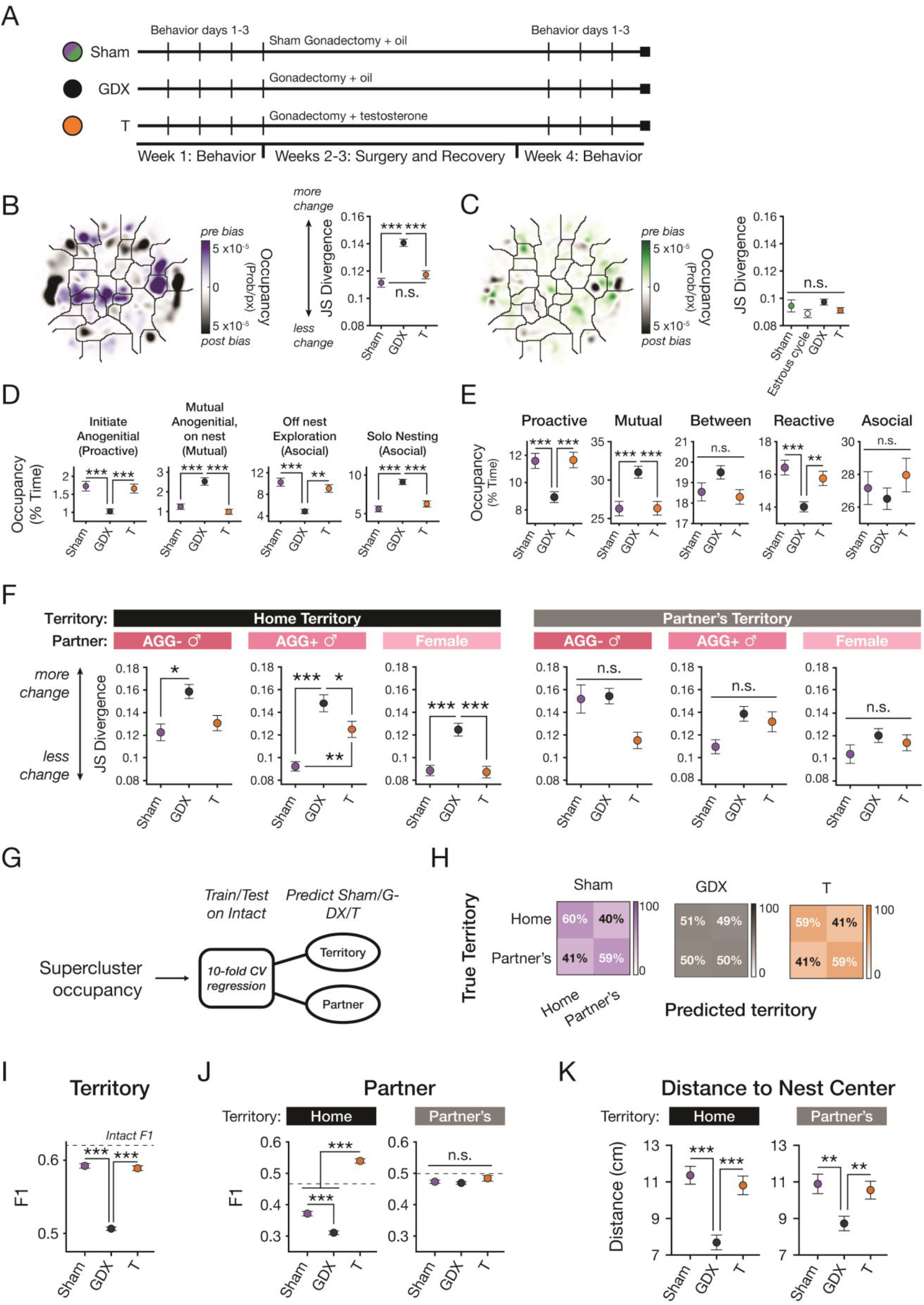
Circulating testosterone is necessary and sufficient for context-specific patterns of behavior in males. A. Experimental timeline. B. Left: Post-GDX - Pre-GDX behavior difference map in males. Right: Jensen-Shannon Divergence from pre- to post-manipulation. N = 432 sham comparisons, 8 mice; N = 756 GDX comparisons, 14 mice; N = 486 T comparisons, 9 mice; α_FDR_ = 0.0333. C. Left: Post-GDX - Pre-GDX behavior difference map in females. Right: Jensen-Shannon Divergence from pre- to post-manipulation or diestrus to estrus. N = 108 sham comparisons, 2 mice; N = 240 estrous comparisons, 20 mice; N = 594 GDX comparisons, 11 mice; N = 648 T comparisons, 12 mice; α_FDR_ = 0.0083. D. Behavior cluster occupancy across hormonal manipulation in males for example proactive, mutual, and asocial behaviors. N = 144 sham sessions, 8 mice; N = 252 GDX sessions, 14 mice; N = 162 T sessions, 9 mice. Initiate anogenital α_FDR_ = 0.0333; Mutual anogenital α_FDR_ = 0.0333; Off-nest exploration α_FDR_ = 0.0333; Solo nesting α_FDR_ = 0.0333. E. Behavior supercluster occupancy across hormonal manipulations. N = 144 sham sessions, 8 mice; N = 252 GDX sessions, 14 mice; N = 162 T sessions, 9 mice. Proactive α_FDR_ = 0.0167; Mutual α_FDR_ = 0.05; Between α_FDR_ = 0.0333; Reactive α_FDR_ = 0.05; Asocial α_FDR_ = 0.05. F. Context-specific Jensen-Shannon Divergence from pre- to post-manipulation in males. Home Territory, AGG-: α_FDR_ = 0.0167; Home Territory, AGG+: α_FDR_ = 0.05; Home Territory, FEM: α_FDR_ = 0.0333; Partner’s Territory, all partners α_FDR_ = 0.05. N_Sham_ = 72 comparisons, 8 mice, N_GDX_ = 126 comparisons, 14 mice, N_T_ = 81 comparisons, 9 mice. G. Decoder schematic. H. Confusion matrices for a territory decoder trained on supercluster occupancy from intact males tested on supercluster occupancy data from post-manipulation (Sham, GDX, or T) males. I. F1 scores for a territory decoder trained on supercluster occupancy from intact males tested on supercluster occupancy data from post-manipulation (Sham, GDX, or T) males. N = 10 folds. α_FDR_ = 0.0333. J. F1 scores for territory-specific partner decoders trained on supercluster occupancy from intact males tested on supercluster occupancy data from post-manipulation (Sham, GDX, or T) males. N = 10 folds. Home Territory α_FDR_ = 0.05; Partner’s Territory α_FDR_ = 0.0167. K. Distance to nest centroid across hormonal manipulations in home and partner’s territory. Both territories: N_Sham_ = 72 sessions, 8 mice; N_GDX_ = 126 behavioral sessions, 14 mice; N_T_ = 81 sessions, 9 mice; α_FDR_ = 0.0333. Data presented as mean ± S.E.M. Linear mixed effects models were used for all statistical tests, unless otherwise stated. Linear Mixed Effects Model formulas can be found in Methods. α threshold set at 0.05, unless otherwise stated. *: *p* < 0.05/α_FDR_, **: *p* < 0.01, ***: *p* < 0.001.

By examining the difference maps from before (pre) to after (post) manipulation in males receiving GDX as a measure of how impact GDX has on behavior, we observe that proactive behaviors are largely pre-GDX biased (Fig. 5B, left), whereas we observe far less behavior change in females receiving GDX (Fig 5C, left). To quantify the extent of behavioral change from pre-to-post manipulation, we used Jensen-Shannon (JS) divergence to compare the probability distributions of fine-grain cluster occupancy before and after the manipulation for each group (Fig. 5B-C). By comparing JS divergence across groups, we find that GDX in males increases JS divergence relative to both the sham and T groups (Fig 5B, right), indicating that T replacement reduces the extent of GDX-induced behavioral change to sham levels. In contrast, examining the pre-to-post-manipulation JS divergence in females reveals no significant differences between any groups or across estrus (Fig 5C, right). This suggests that broad patterns of social behavior in males are more sensitive to changes in hormone profile, potentially due to differences in the initial hormonal milieu and circuit architecture.

To better understand which specific behaviors are impacted by GDX, we compared behavioral occupancy from all partner-territory contexts across manipulation groups using both our fine- (clusters) and coarse-grain (superclusters) behavioral strategies (Fig. 5D-E; Fig. S7B-F). At the level of individual behavior clusters, GDX induces behavioral change in a variety of behavioral clusters rather than a subset of specific social actions. For instance, GDX increases both social and asocial behavior on the nest while reducing off-nest behaviors and decreasing the average distance between the subject mouse and the nest center in both the home and partner’s territory (Fig. 5D, K; Fig S7B-F). When we examine behavioral occupancy at a coarse-grain level, we find that GDX reduces both proactive and reactive behavior while increasing mutual social interaction (Fig. 5E). When we examine behavioral persistence, or mean time spent engaged in each supercluster bout, we find that GDX reduces proactive behavior persistence and increases the time spent engaged in each mutual interaction (Fig. S7A), consistent with the effect of GDX on behavioral occupancy. However, there is no effect of GDX on reactive behavior persistence (Fig. S7A). Combined with the reduced reactive occupancy, these data reveal that mice engage in fewer overall bouts of reactive behaviors following GDX.

We find that GDX effects on behavior are context specific. Using JS divergence to compare behavior from before to after hormonal manipulation, we find GDX significantly affects behavior only in the home territory, across partners. T replacement promotes a full or partial behavioral rescue in each of these contexts (Fig. 5F). Finally, we compared patterns of behavior by training a binomial logistic regression on pre-manipulation supercluster occupancy to decode territory or partner from post-manipulation behavior (Fig. 5G). We find that, compared to sham and T groups, GDX significantly decreases territory decoding accuracy (Fig. 5H-I). Strikingly, GDX significantly decreases partner decoding accuracy only in the home territory (Fig. 5J). Together, these data indicate that the GDX primarily impacts behavioral action in the home territory. With one exception (Cluster 14; Fig. S7E), T replacement fully or partially rescues all the observed GDX-induced behavioral changes in males irrespective of whether we take a fine-or coarse-grained approach to quantify behavior (Fig. 5; Fig. S7).

Overall, these results demonstrate that circulating gonadal hormones in males are required for high levels of proactive behavior and that exogenous T can rescue these behavioral changes. Further, they show that GDX-induced behavioral change occurs primarily in the home territory and affects behavioral action beyond simple aggression. This suggests that T does not simply generate proactive behavior but enables context-specific patterns of behavior.

### Loss of circulating gonadal hormones disrupts context rescaling

Finally, to test the necessity of circulating gonadal hormones for context-dependent rescaling of neural activity, we recorded neural activity in males who had undergone either GDX or sham GDX (surgery but no gonadectomy). By comparing neural activity between these two groups across the 6 social contexts (Fig. 6A), we observe sweeping differences that span behaviors and contexts (Fig. 6A-C). Notably, differences in behavior-specific response across contexts are not unidirectional: During interactions with males, GDX decreases neural activity, whereas during interactions with females, GDX increases neural activity relative to sham surgery (Fig. 6B). By computing the cosine distance between vectors of mean neural population activity across the fine-grain behavioral clusters, we can determine the effect of these changes on network code similarity. We find GDX, relative to sham surgery, reduces cosine distance between FEM and AGG+ social action codes in both territories and increases the cosine distance between AGG+ and AGG-males in the mouse’s home territory (Fig. 6C). Taken together, these data suggest that loss of circulating hormones severely disrupts the context-specific encoding of behavioral action that we observe in gonadally intact animals.

**Figure 6.**
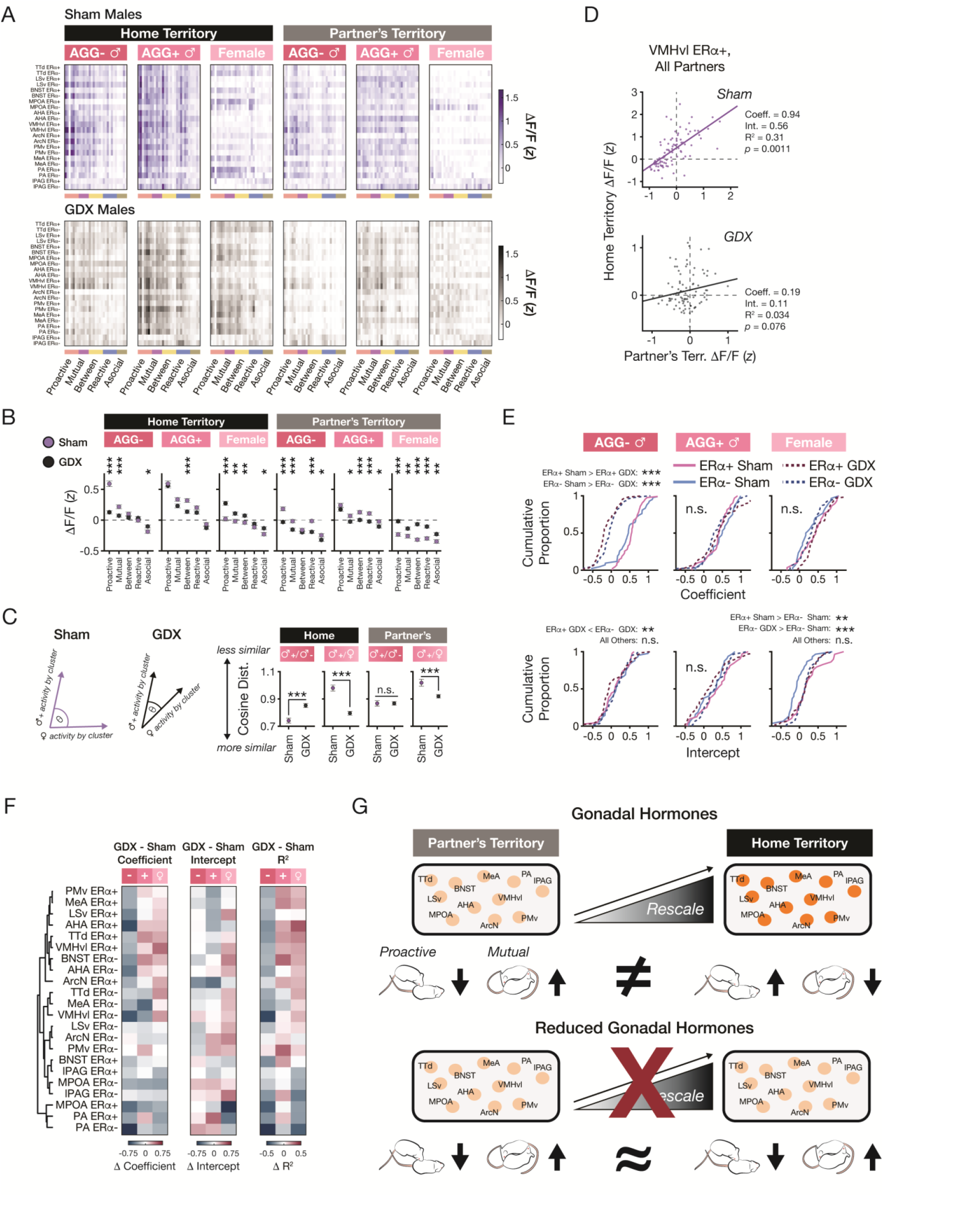
Social action code rescaling requires circulating gonadal hormones. A. Heatmaps of mean neural activity across all recorded populations and behavior clusters in each social context. Top: Sham male subjects. Bottom: GDX male subjects. B. Mean supercluster-associated neural population activity as a function across all 22 SBN populations in sham (purple) or GDX (black) male subject mice. Home Territory, AGG-: α_FDR_ = 0.03, N_Sham_ = 260 populations, N_GDX_ = 387 populations; Home Territory, AGG+: α_FDR_ = 0.01, N_Sham_ = 238 populations, N_GDX_ = 387 populations; Home Territory, FEM: α_FDR_ = 0.04, N_Sham_ = 238 populations, N_GDX_ = 387 populations; Partner’s Territory, AGG-: α_FDR_ = 0.04, N_Sham_ = 259 populations, N_GDX_ = 387 populations; Partner’s Territory, AGG-: α_FDR_ = 0.04, N_Sham_ = 233 populations, N_GDX_ = 388 populations; Partner’s Territory, FEM: α_FDR_ = 0.05. N = 4 sham mice, 6 GDX mice, 3 sessions per mouse. C. Cosine distance between social action code in sham (purple) or GDX (black) male subject mice during interactions with AGG+ and AGG-partners or AGG+ and FEM partners. Home Territory, AGG+ vs AGG-: *p* = 9.30 × 10^−6^, N_Sham_ = 345 within-population comparisons, 4 mice, N_GDX_ = 770 within-population comparisons, 6 mice; Home Territory, AGG+ vs FEM: *p* =5.81 × 10^−14^, N_Sham_ = 345 within-population comparisons, 4 mice, N_GDX_ = 770 within-population comparisons, 6 mice; Partner’s Territory, AGG+ vs AGG-: *p* = 0.99, N_Sham_ = 345 within-population comparisons, 4 mice, N_GDX_ = 770 within-population comparisons, 6 mice; Partner’s Territory, AGG+ vs FEM: *p* = 1.30 × 10^−4^, N_Sham_ = 345 within-population comparisons, 4 mice, N_GDX_ = 770 within-population comparisons, 6 mice. D. Example cross-territory regression from the VMHvl ERα+ population of a representative sham (top) and GDX (bottom) male subject mouse. Each data point represents the z-scored neural activity in the home and partner’s territories for a given partner-behavior cluster pairing. E. Distributions of regression coefficients (top) and intercepts (male) in sham and GDX subject male mice, separated by interaction partner. Statistical significance determined with Kolmogorov-Smirnov tests. AGG-Partner, Coefficient: α_FDR_ = 0.025, Intercept: α_FDR_ = 0.0125, N_Sham_ = 43 ERα+ populations, 42 ERα+ populations, 4 mice, N_GDX_ = 63 ERα+ populations, 62 ERα+ populations, 6 mice. AGG+ Partner, Coefficient: α_FDR_ = 0.05, Intercept: α_FDR_ = 0.0125, N_Sham_ = 32 ERα+ populations, 32 ERα+ populations, 4 mice, N_GDX_ = 64 ERα+ populations, 60 ERα+ populations, 6 mice. FEM Partner, Coefficient: α_FDR_ = 0.05, Intercept: α_FDR_ = 0.025, N_Sham_ = 29 ERα+ populations, 30 ERα+ populations, 4 mice, N_GDX_ = 63 ERα+ populations, 62 ERα+ populations, 6 mice. F. Heatmaps of the difference between GDX and Sham regression coefficients, intercepts, and R^2^ values (left to right) as a function of SBN population. SBN populations are hierarchically clustered based on coefficient and intercept differences. G. Summary of findings. Data presented as mean ± S.E.M. Linear mixed effects models were used for all statistical tests, unless otherwise stated. Linear Mixed Effects Model formulas can be found in Methods. α threshold set at 0.05, unless otherwise stated. *: *p* < 0.05/α_FDR_, **: *p* < 0.01, ***: *p* < 0.001.

To further quantify this, we used a linear model to quantify territory-dependent rescaling of the neural action code and compared model coefficients and intercepts on a per signal basis. Similar to the intact animals, we fit each signal with a simple linear model to predict home territory behavior-associated neural population response based on the neural population response during the same behaviors performed in the partner’s territory (Fig. 6D).

Whether using data from all partners (Fig. 6D) or individual partner identities (*e*.*g*., AGG-males; Fig. 6E-F), we observe differences in the model fit and parameters between the GDX and sham individuals, on a per-signal basis and across multiple populations. At the level of the entire SBN, we observe partner-specific differences in the coefficients and intercepts between GDX and sham GDX males. Specifically, the greatest difference in fit coefficient occurs during interactions with AGG-males, and the largest differences in the intercept occur during interactions with females. Next, we used the differences in coefficient and intercept to cluster SBN neural populations. With this analysis, we find three major subnetworks with unique patterns of GDX-induced change (Fig. 6F). We observe a subnetwork of largely ERα+ neural populations undergoes an inverse shift in the responses, with a decrease in rescaling during AGG-interactions and an increase during AGG+ and FEM interactions. Additionally, we observe a second subnetwork of largely ERα− neural populations that undergo a decrease in rescaling regardless of partner type. Finally, we observe a small subnetwork consisting of both ERα+ and ERα− PA populations and the ERα+ MPOA population which shows a unique decrease in rescaling and intercept during FEM interactions that does not occur in the other subnetworks. These data suggest that these subnetworks may share independent sources of noise but are yoked together in the presence of circulating hormones.

Overall, these data underscore the broad control that circulating gonadal hormones, particularly testosterone, have on context-specific behavioral action and its network-scale encoding (Fig. 6G). This suggests a new model or hormonal function whereby testosterone acts to tune SBN-wide neural activity in a territory-dependent manner. Therefore, gonadal hormones therefore enable a mapping of social context to patterns of context-specific behavior.

## Discussion

Here, we demonstrate for the first time that social context, in particular territory, naturally and rapidly rescales the SBN action code, boosting behavior-associated activity during social interactions at home. Further, we show that both ERα+ and ERα− populations in the SBN encode social context-specific patterns of behavioral action in a distributed manner. These context-specific patterns of social behavior were sensitive to loss of circulating gonadal hormones via GDX in adulthood in males but not females, and T replacement following GDX was sufficient to rescue these behavioral changes. These behavioral changes were most prominent in the home territory of the mice and were associated with an SBN-wide disruption of behavior encoding and territorial rescaling of the neural code. Strikingly, GDX-induced changes to territorial rescaling differed between SBN ERα+ and ERα− populations, suggesting unique sources of input to these populations and potential functional divergence following loss of circulating hormones.

We took a comprehensive and fully unsupervised approach to identify and quantify all behavior at a frame-by-frame level, during social interaction between two mice, allowing us to capture hierarchical behavioral organization consistent with previous work^35^ and to discover that the social context rescales the entire behavior code. While it has previously been shown that social context gates aggressive behavior^2^, we found that social context modifies the behavioral repertoire across the entirety of the behavioral landscape rather than influencing a handful of specific aggressive behaviors. Consistent with a role for the SBN in encoding the entirety of the behavioral landscape, we found that the social action code rescales, even when proactive behaviors, such as aggression, are not included in this analysis.

Our data shows that social context enables a rapid adaptation of a social action code, suggesting a fast, hormone-dependent rescaling of social sensory input. Rapid adaptation of neural response magnitudes to the same input or action across contexts is a canonical feature of sensory and motor circuits and occurs on the timescale of seconds^43–46^. One possibility is that rapid brain-wide rescaling emerges due to increased hormone-mediated intra-SBN recurrent activity, which could propagate shared signals throughout the network^47,48^. For example, circulating gonadal hormones promote synaptic plasticity within the SBN, including both increasing dendritic spine density and potentiating synaptic responses to afferents^49–52^. Indeed, *in silico* models of recurrent neural networks have demonstrated that both local recurrency as well as afferent activity from other brain regions is required for the propagation of neural activity^53^.

In the social behavior network of adult mice, gonadal hormones acting through ERα have been shown to drive the transcription of genes associated with neural excitability, circuit wiring, and synaptic plasticity genes^54,55^, raising the intriguing possibility that circulating T promotes recurrent signaling between the ERα+ and ERα− subnetworks to enable territorial rescaling of neural responses. In support of this ability, previous work has demonstrated the potential for circuit-wide remodeling by endogenous and exogenous gonadal hormones^49,51^. In addition, circulating gonadal hormones influence the magnitude of neural responses to social sensory cues in primary sensory neurons^56^, sensory cortex^57^, and the SBN^22,58^. As such, the GDX-induced impairment of territorial rescaling of social action could result from lost upstream social sensory input, lost intra-SBN recurrent activity, or both. Future computational neuroendocrinology experiments will elucidate the key neural pathways that enable territorial rescaling.

Acute responses to gonadal hormones themselves might also mediate context rescaling. In addition to genomic effects on hormone receptor expression neurons, T and T-derived estrogen can directly change neural activity through GPCRs as well as the membrane associated ion channel TRPM8^59,60^. In male rodents, T is released acutely during social interaction and in a partner specific manner with higher levels of T release occurring during interactions with females in estrus^61,62^. This suggests that T could acutely update the SBN action code to match an individual’s behavior to its social context. As pulsatile T release occurs in males and not females, this would predict a sex-specific impact of GDX on context-dependent patterns of behavioral action Indeed, we found that GDX uniquely disrupts the behavioral landscape of males but not females. Moreover, we found that GDX promotes an “away-like” behavioral landscape, including lower levels of proactive behavior and higher levels of mutual behaviors. In addition, we observe a decrease in the frequency of reactive behaviors, which may reflect less engagement from the partner mouse due to behavioral or pheromonal change post-GDX^63,64^. While it has been previously shown that GDX in male mice reduces aggression regardless of strain, the magnitude of this reduction is strain-specific^39^. This suggests that, while T’s role in promoting proactive behaviors is likely conserved across species and strains^6,7,39,65,66^, differences in hormone receptor expression patterns may constrain its role.

Importantly, we observed that hormonal modulation of behavior goes beyond social interaction. Tracking the location of the nest allowed us to observe additional differences in specific non-social behaviors across contexts and hormone states. For example, GDX reduced off-nest behaviors in male mice relative to sham controls and mice treated with T. This T-dependent bias for off-nest behavior is consistent with prior work demonstrating T increases goal-driven exploration of a local environment^67^. Combined with our results demonstrating a T-dependent reduction in proactive behaviors, these results support a model whereby T increases motivation for and persistence of goal-directed behaviors, whether they be social or exploratory.

In contrast to the large effects of GDX on male behavior, we found that neither GDX, T replacement, nor estrous stage significantly impacted the behavioral landscape during social interaction in female mice. These data complement recent findings demonstrating that estrous stage does not drive variability in the landscape of spontaneous solo behavior in females^68^. Nevertheless, estrous stage and GDX do influence the display of specific social actions in females, such as opposite-sex approach^22^ and mating behaviors^49,52,58^. Taken together, these data suggest that circulating gonadal hormones in female mice impact specific behavioral actions rather than the totality of the behavioral landscape as seen in males.

Our finding that territorial rescaling of SBN activity is disrupted when hormone levels are low suggests that a mouse’s ability to identify both its partner and the location of their interaction may be impaired, particularly in the home territory. Several lines of evidence from our neural data support this idea. First, we observe that disruptions in scaling do not always reduce activity. Post-GDX, we find a complete flattening of the behavior tuning curve and loss of territorial rescaling across SBN populations in males during interactions with less aggressive male partners. In contrast, we find widespread disinhibition of neural activity in males during interactions with female partners. This disinhibition results in a more male partner-like SBN neural code irrespective of territory. Second, our recordings identified a unique subnetwork of ERα+ and ERα− PA and ERα+ MPOA populations that undergo a GDX-induced reduction in both territory regression coefficient and intercept for female partners. These data are consistent with neural activity misreading the identity of female partners regardless of social territory. In combination with the neural activity changes occurring during interactions with less aggressive males, this could promote confusion in identifying the social context.

This subnetwork supports previous work that identifies a mating-biased SBN from recordings of ERα+ SBN populations^13^ in male mice, and extends this finding to ERα− populations and to female mice, suggesting that ERα+ may not represent a specialized population. While some previous studies have suggested that ERα− populations do not play a causal role in in the generation of specific social behaviors^12,19^, we find that ERα− populations robustly encode a broad variety of social behaviors and are part of the distributed neural code for behavior, partner, and territory in both sexes. Further, by comparing neural activity across contexts, we discovered territory’s role in rescaling the social action code in both SBN ERα+ and ERα− populations and that GDX disrupted territorial rescaling in both ERα+ and ERα− neurons in a population-specific manner. These findings are consistent with sequencing studies demonstrating the expression of non-ERα hormone receptors (e.g., ERβ, androgen receptor) in both ERα+ and ERα− populations throughout the SBN^54,55,69,70^.

Taken together, our data uncover a key, hormone-sensitive role for both SBN ERα+ and ERα− populations in the encoding of social action and related contextual variables.

Though decades of research have suggested the role of gonadal hormones in territory establishment and defense^23–29^, direct evidence for how it exerts this modulation has been lacking. Here, we provide the first demonstration of hormonal control of large-scale neural activity, showing that social context-specific patterns of behavioral action co-occur with a network-level rescaling of the SBN behavioral action code. Further, using a hormonal perturbation and rescue approach, we show that both the context-specific patterns of behavior and neural activity rescaling are sensitive to a loss of circulating gonadal hormones. While many have hypothesized that circulating gonadal hormones influence behavior by coordinating SBN neural population activity dynamics^6–10^, our data provide, for the first time, a mechanistic explanation of how circulating hormones influence neural populations at the network-scale to generate context-specific patterns of behavioral action.

## Author Contributions

Conceptualization: E.M.G., A.L.F. Data curation: E.M.G., L.S., A.L. Formal analysis: E.M.G. Funding acquisition: E.M.G., A.L.F. Investigation: E.M.G., L.S. Methodology: E.M.G., J.I.G. Resources: E.M.G., A.L.F. Software: E.M.G., J.I.G., L.S. Supervision: A.L.F. Validation: E.M.G., J.I.G., A.L. Visualization: E.M.G. Writing – original draft: E.M.G., A.L.F. Writing - review & editing: E.M.G., J.I.G., A.L.F.

## Data Availability

All behavior and neural data will be made publicly available on and source data for all figures will be available following publication of this manuscript.

## Code Availability

All code will be made available on GitHub repository by date of publication: https://github.com/FalknerLab/ContextRescaling and https://github.com/emguthman/ContextRescaling.

## Acknowledgments

We thank M. Murthy, S. Correa, E. van Veen, M. Anderman, and members of the Falkner laboratory for useful discussions; K. Aghi, I. Alcantara, D. Allen, M. Asokan, A. Bondy, S. Sun for feedback on the manuscript; S. Oline for feedback on and assistance with hardware engineering; and, T. Yamaguchi for sharing the Cre-excluding GCaMP6f virus. Funding was from NIH K99MH135212 (to E.M.G.), NIH F32MH126562 (to E.M.G.), DP2MH126375 (to A.L.F.), NIH R01MH126035 (to A.L.F.), NYSCF (to A.L.F.), Simons Foundation(SCGB) (to A.L.F.), Klingenstein Foundation (to A.L.F.), and Alfred P. Sloan Fellowship (to A.L.F.), McKnight Foundation (A.L.F), Allen Institute (A.L.F). A.L.F. is a New York Stem Cell Foundation Robertson Investigator.

## Declaration of Interests

E.M.G. is a co-founder of the 501(c)3 nonprofit organization Community Estrogen, Inc. and is a member of its board of directors. E.M.G. receives no financial compensation from Community Estrogen, Inc. Authors declare no other competing interests.

**Figure S1.**
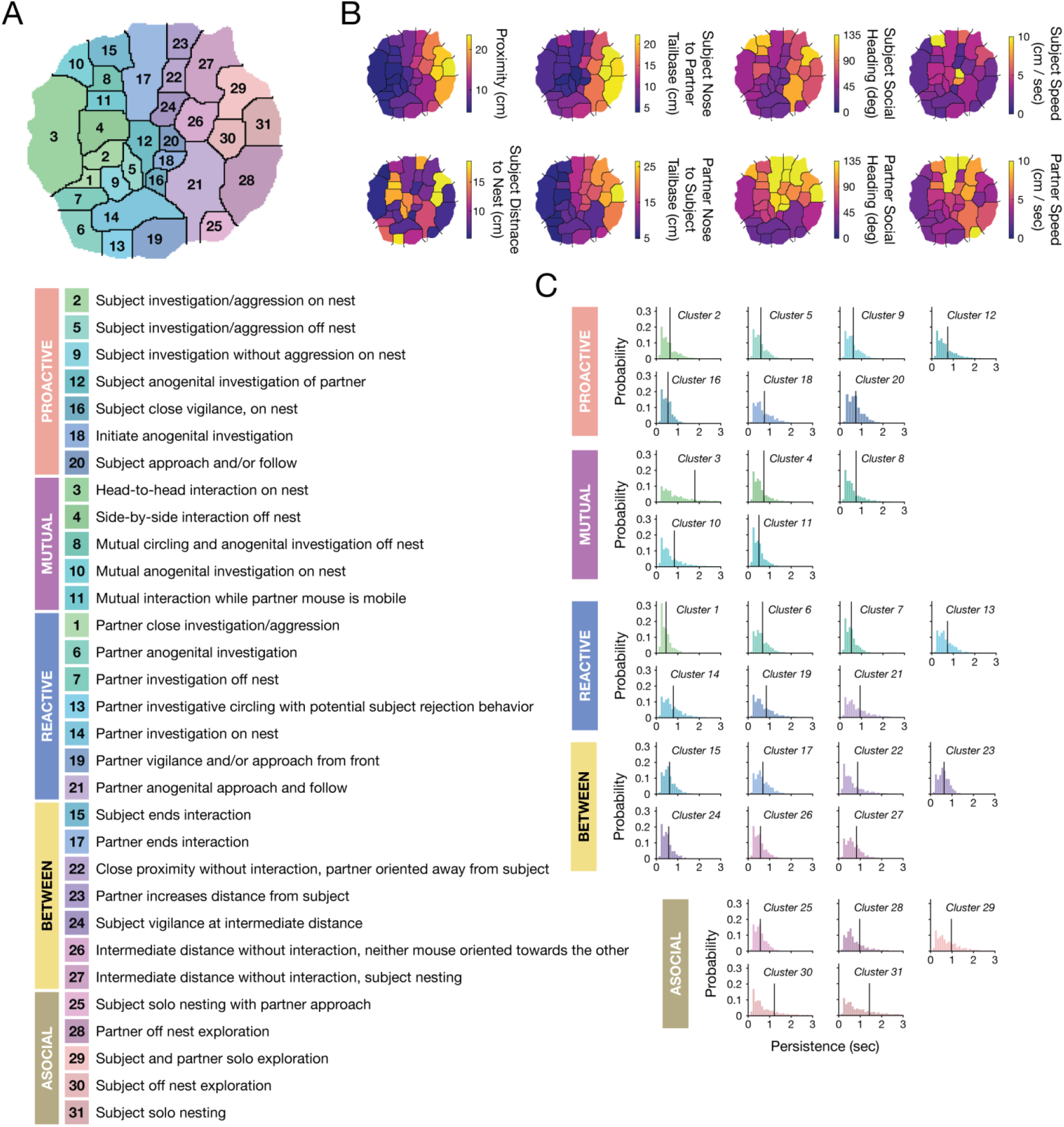
Behavior cluster characterization. A. Cluster identities. Clusters ordered by intermouse proximity and grouped by Supercluster. B. Mean feature values plotted onto behavior maps for example features. C. Cluster persistence histograms. Black line designates the mean value. X limits cropped to 0 to 3 seconds for display purposes.

**Figure S2.**
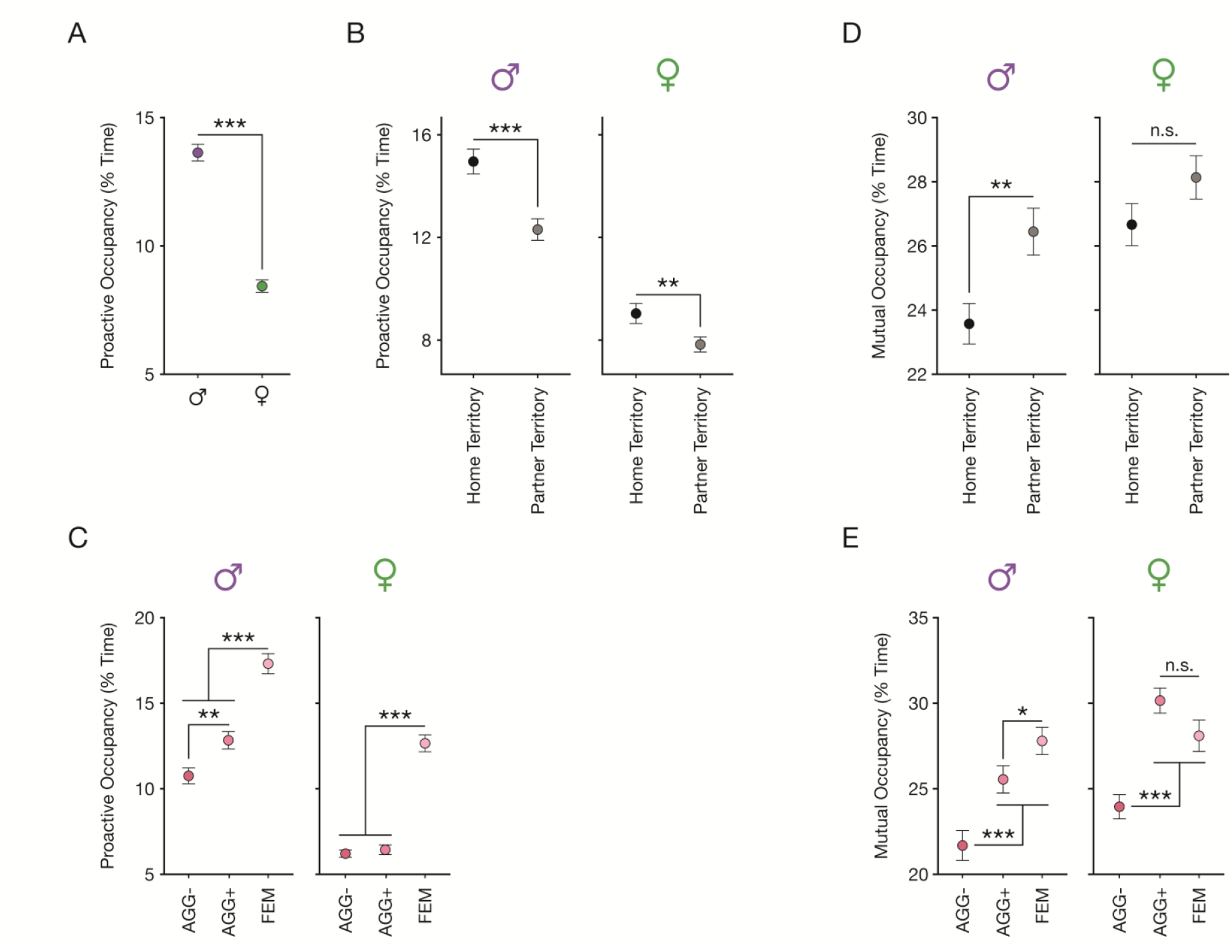
Sex and social context determine levels of proactive behavior. A. Proactive behavior in male and female subject mice. Linear Mixed Effects Model, *p =* 4.97 × 10^−16^, N = 558 male subject sessions, 31 mice, 486 female subject sessions, 27 mice. B. Proactive behavior in home and partner territories for male and female subject mice. Linear mixed effects models; Males: *p =* 3.81 × 10^−6^, N = 279 sessions per territory, 31 mice; Females: *p =* 0.0013, 243 sessions per territory, 27 mice. C. Proactive behavior during male and female subject interactions with AGG-, AGG+, and FEM partners. Linear mixed effects models; Males: α_FDR_ *=* 0.05, N = 186 sessions per partner, 31 mice; Females: α_FDR_ *=* 0.0333, 162 sessions per partner, 27 mice. D. Mutual behavior in home and partner territories for male and female subject mice. Linear mixed effects models; Males: *p =* 0.0014, N = 279 sessions per territory, 31 mice; Females: *p =* 0.098, 243 sessions per territory, 27 mice. E. Proactive behavior during male and female subject interactions with AGG-, AGG+, and FEM partners. Linear mixed effects models; Males: α_FDR_ *=* 0.05, N = 186 sessions per partner, 31 mice; Females: α_FDR_ *=* 0.0333, 162 sessions per partner, 27 mice. Data presented as mean ± S.E.M. Linear Mixed Effects Model formulas can be found in Methods. *: *p* < α_FDR_, **: *p* < 0.01, ***: *p* < 0.001.

**Figure S3.**
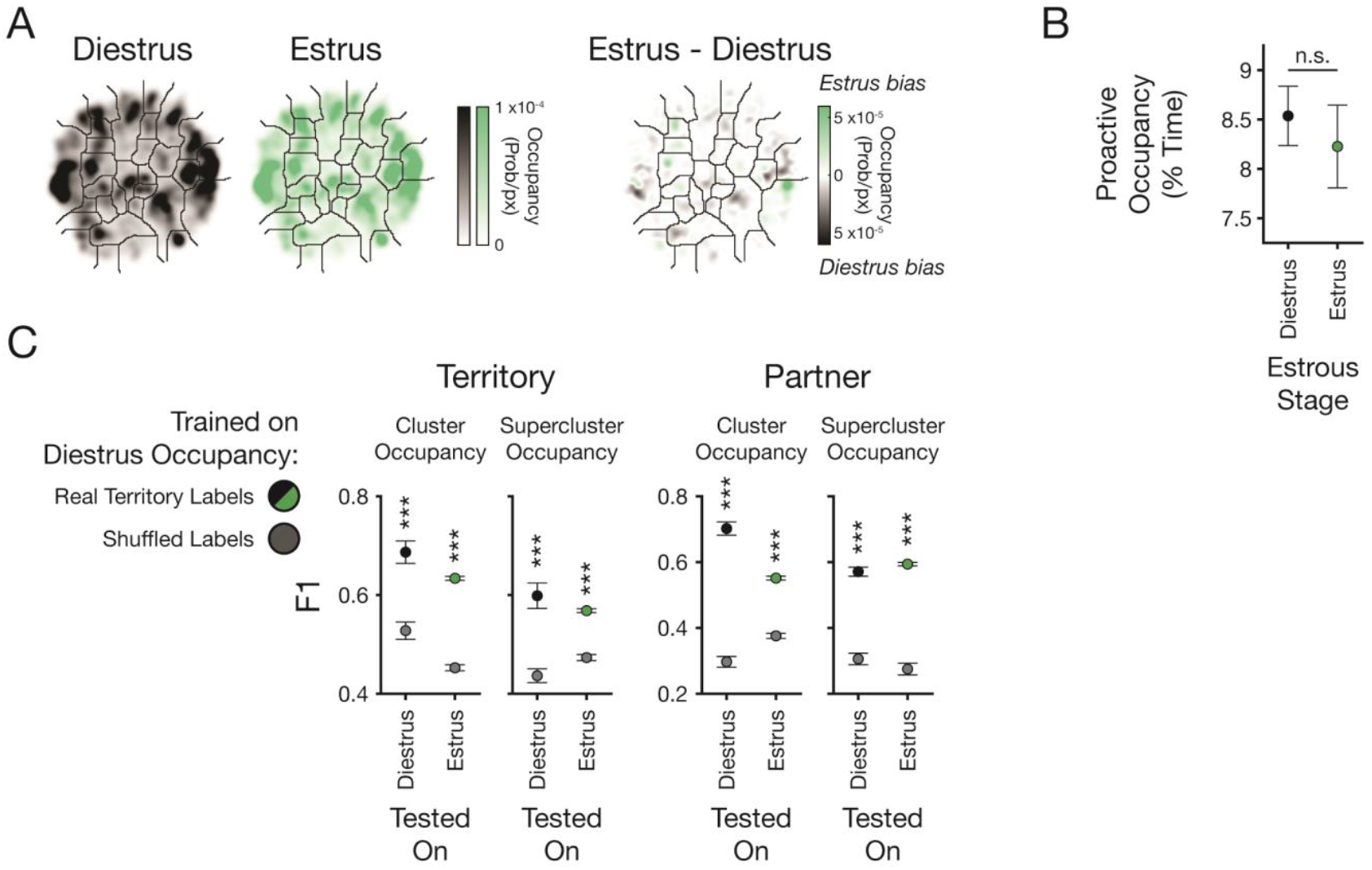
Patterns of behavior during social interaction do not change across estrous stages. A. Left: Mean behavior maps from females in estrus or diestrus. Right: Estrus - Diestrus difference map. B. Proactive behavior occupancy in female subjects during estrus or diestrus. Linear mixed effects model, *p* = 0.62, N = 162 estrus sessions, 21 mice, 324 diestrus sessions, 26 mice. C. Results of 10-fold CV linear territory (left) and partner (right) decoders trained on diestrus cluster or supercluster occupancies and tested on diestrus or estrus occupancies. Territory, cluster: *p*_Diestrus_ = 2.19 × 10^−4^, *p*_Estrus_ = 2.93 × 10^−10^, paired t-tests, N = 10 folds; Territory, supercluster: *p*_Diestrus_ = 6.06 × 10^−4^, *p*_Estrus_ = 4.99 × 10^−8^, paired t-tests, N = 10 folds; Partner, cluster: *p*_Diestrus_ = 1.43 × 10^−7^, *p*_Estrus_ = 2.82 × 10^−9^, paired t-tests, N = 10 folds; Partner, supercluster: *p*_Diestrus_ = 2.72 × 10^−6^, *p*_Estrus_ = 1.98 × 10^−8^, paired t-tests, N = 10 folds. Data presented as mean ± S.E.M. Linear Mixed Effects Model formulas can be found in Methods. *: *p* < 0.05, **: *p* < 0.01, ***: *p* < 0.001.

**Figure S4.**
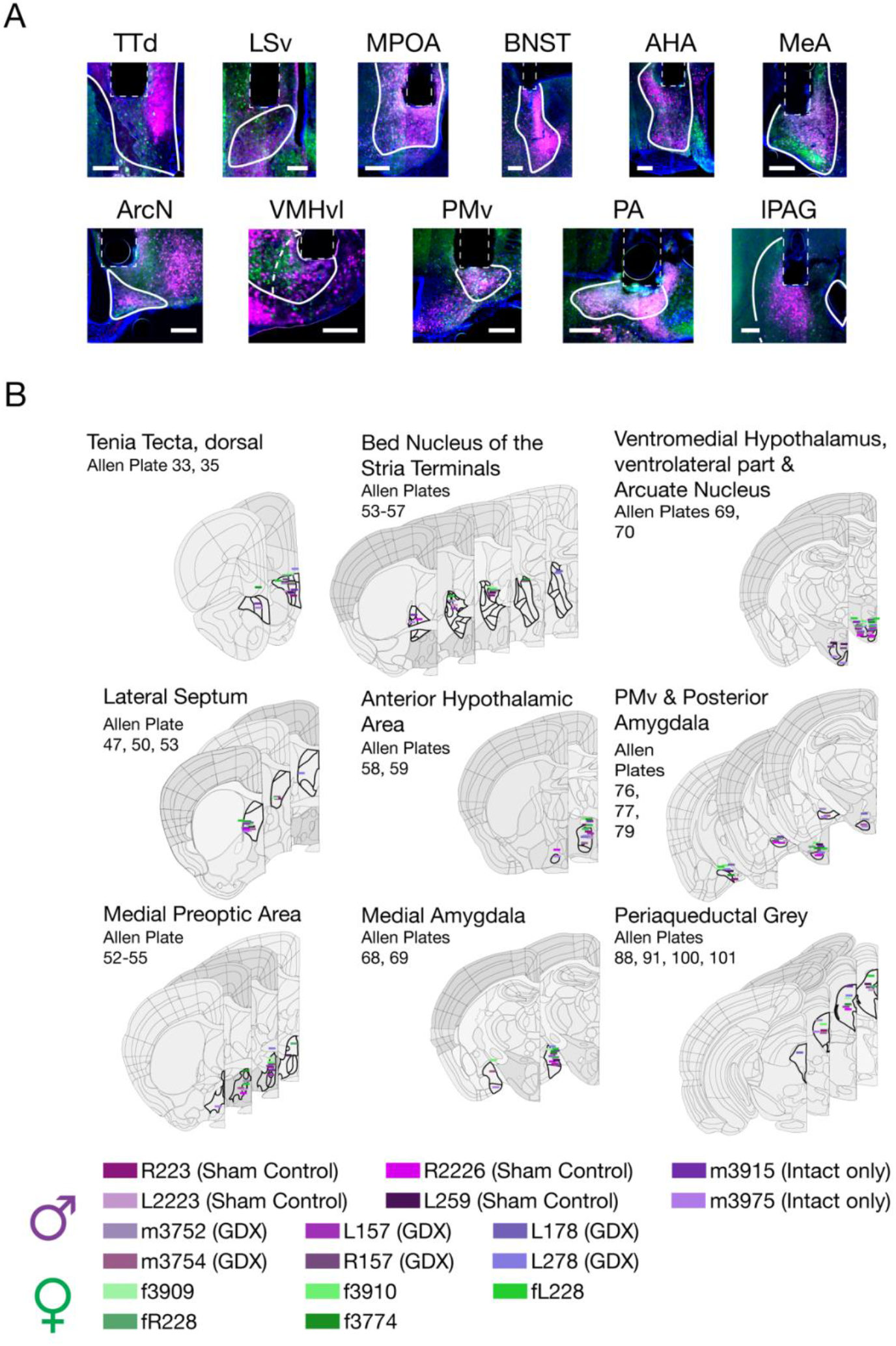
Multisite photometry histology. A. Example viral and fiber targeting. DAPI displayed in blue, GCaMP6f in green, and jRCaMP1b in magenta. B. Location of all fibers in all imaging mice.

**Figure S5.**
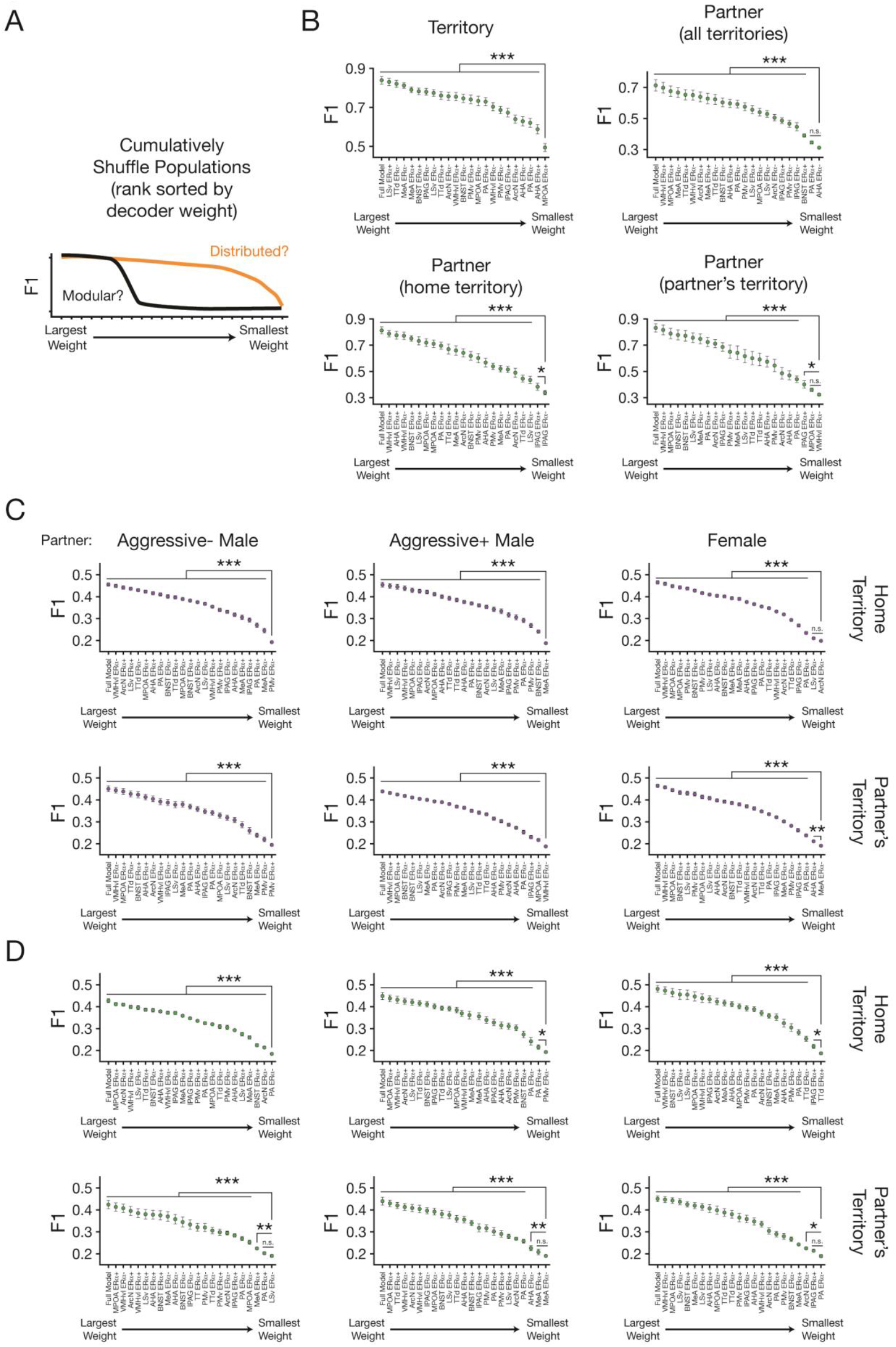
Distributed encoding of social context and action by the social behavior network encodes. A. Schematic depicting modular vs. distributed outcomes. B. Cumulatively shuffled 10-fold CV territory and partner decoders in female mice. Linear mixed effects model used to compare decoding in the fully shuffled condition (i.e. all populations from largest through smallest weight shuffled) against all other runs of the cumulatively decoder. Territory decoding: MPOA ERα+ vs. all other pops, all *p* < 5.94 × 10^−8^. Partner decoding (all territories): AHA ERα− vs. PA ERα+, *p* = 0.08; AHA ERα− vs. all other pops, all *p* < 8.98 × 10^−5^. Partner decoding (home territory): lPAG ERα− vs. lPAG ERα+, *p* = 0.032; lPAG ERα− vs. all other pops, all *p* < 1.061 × 10^−5^. Partner decoding (partner’s territory): VMHvl ERα− vs. MPOA ERα−, *p* = 0.22; VMHvl ERα− vs. lPAG ERα+, *p* = 0.011; VMHvl ERα− vs. all other pops, all *p* < 1.45 × 10^−4^. C. Same as A, but for context-specific action decoders in males. Home territory, AGG-: PMv ERα− vs. all other pops, all *p* < 1.17 × 10^−12^. Home territory, AGG+: MeA ERα+ vs. all other pops, all *p* < 3.27 × 10^−11^. Home territory, FEM: ArcN ERα− vs. LSv ERα−, *p* = 0.10; ArcN ERα− vs. all other pops, all *p* < 8.71 × 10^−7^. Partner’s territory, AGG-: PMv ERα+ vs. all other pops, all *p* < 1.78 × 10^−4^. Partner’s territory, AGG+: PMv ERα− vs. all other pops, all *p* < 3.00 × 10^−6^. Partner’s territory, FEM: MeA ERα− vs. AHA ERα+, *p* = 0.0017; MeA ERα− vs. all other pops, all *p* < 2.23 × 10^−12^. D. Same as A, but for context-specific action decoders in females. Home territory, AGG-: PA ERα− vs. all other pops, all *p* < 3.97 × 10^−4^. Home territory, AGG+: PMv ERα− vs. PA ERα+, *p* = 0.012; PMv ERα− vs. all other pops, all *p* < 3.64 × 10^−7^. Home territory, FEM: TTd ERα+ vs. lPAG ERα+, *p* = 0.011; TTd ERα+ vs. all other pops, all *p* < 4.03 × 10^−7^. Partner’s territory, AGG-: LSv ERα− vs. PA ERα+, *p* = 0.29; LSv ERα− vs. MeA ERα+, *p* = 0.0037; LSv ERα− vs. all other pops, all *p* < 4.15 × 10^−7^. Partner’s territory, AGG+: MeA ERα− vs. MeA ERα+, *p* = 0.095; MeA ERα− vs. AHA ERα−, *p* = 0.0013; MeA ERα− vs. all other pops, all *p* < 1.00 × 10^−8^. Partner’s territory, FEM: PA ERα− vs. lPAG ERα+, *p* = 0.090; PA ERα− vs. ArcN ERα−, *p* = 0.014; MeA ERα− vs. all other pops, all *p* < 3.44 × 10^−4^. N = 12 male models, 5 female models. Linear Mixed Effects Model formulas can be found in Methods. Data presented as mean ± S.E.M. *: *p* < 0.05, **: *p* < 0.01, ***: *p* < 0.001.

**Figure S6.**
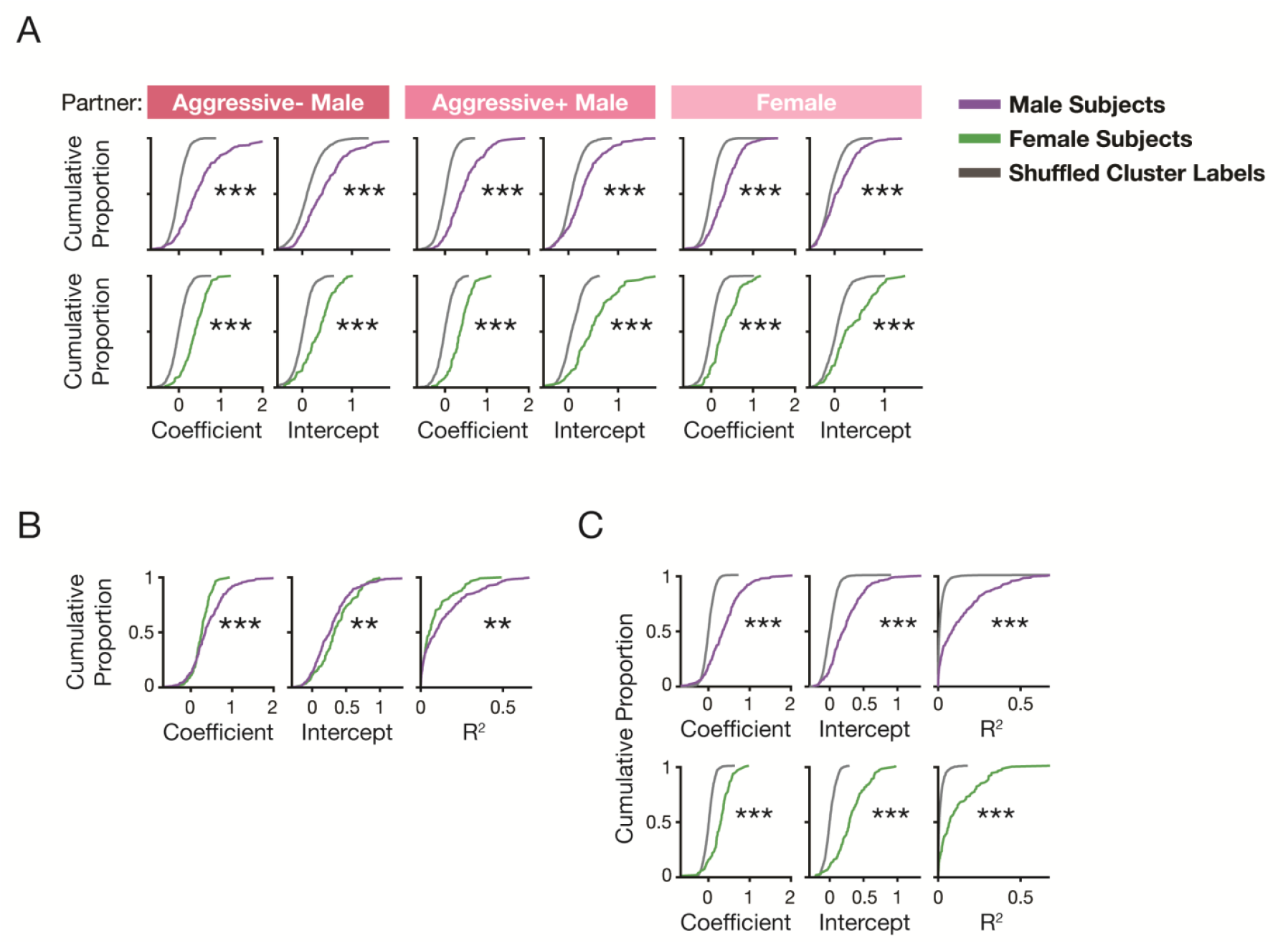
Territorial gain control is robust across partners and behaviors. A. Left to right: Distributions of partner-specific regression coefficients, intercepts, and R^2^ in male (top) and female (bottom) subjects compared to shuffle (Male, AGG-: *p*_Coefficient_ = 6.00 × 10^−66^, *p*_Intercept_ = 4.21 × 10^−27^, *p*_R-Squared_ = 3.41 × 10^−44^, N = 252 populations, 12 mice; Male, AGG+: *p*_Coefficient_ = 3.96 × 10^−65^, *p*_Intercept_ = 7.07 × 10^−25^, *p*_R-Squared_ = 4.74 × 10^−39^, N = 252 populations, 12 mice; Male, FEM: *p*_Coefficient_ = 1.43 × 10^−46^, *p*_Intercept_ = 7.89 × 10^−15^, *p*_R-Squared_ = 7.57 × 10^−30^, N = 251 populations, 12 mice; Female, AGG-: *p*_Coefficient_ = 4.73 × 10^−34^, *p*_Intercept_ = 1.53 × 10^−25^, *p*_R-Squared_ = 2.13 × 10^−23^, N = 106 populations, 5 mice; Female, AGG+: *p*_Coefficient_ = 1.94 × 10^−34^, *p*_Intercept_ = 1.06 × 10^−24^, *p*_R-Squared_ = 1.33 × 10^−22^, N = 107 populations, 5 mice; Female, FEM: *p*_Coefficient_ = 4.47 × 10^−22^, *p*_Intercept_ = 8.61 × 10^−14^, *p*_R-Squared_ = 1.06 × 10^−14^, N = 106 populations, 5 mice; Kolmogorov-Smirnov tests). B. Comparison of regression coefficients, intercepts, and R^2^ between male and female subject mice (Kolmogorov-Smirnov tests; *p*_Coefficient_ = 2.86 × 10^−5^, *p*_Intercept_ = 0.0088, *p*_R-Squared_ = 0.0042; N_Male_ = 257 populations, 12 mice, N_Female_ = 107 populations, 5 mice). C. Cross-territory regression coefficients, intercepts, and R^2^, computed without proactive clusters, in male (top) and female (bottom) subject mice compared to shuffle (Male: *p*_Coefficient_ = 3.51 × 10^−71^, *p*_Intercept_ = 1.05 × 10^−53^, *p*_R-Squared_ = 7.28 × 10^−58^, N = 257 populations, 12 mice; Female: *p*_Coefficient_ = 9.12 × 10^−38^, *p*_Intercept_ = 6.17 × 10^−46^, *p*_R-Squared_ = 1.30 × 10^−27^, N = 107 populations, 5 mice; Kolmogorov-Smirnov tests). *: *p* < 0.05, **: *p* < 0.01, ***: *p* < 0.001.

**Figure S7.**
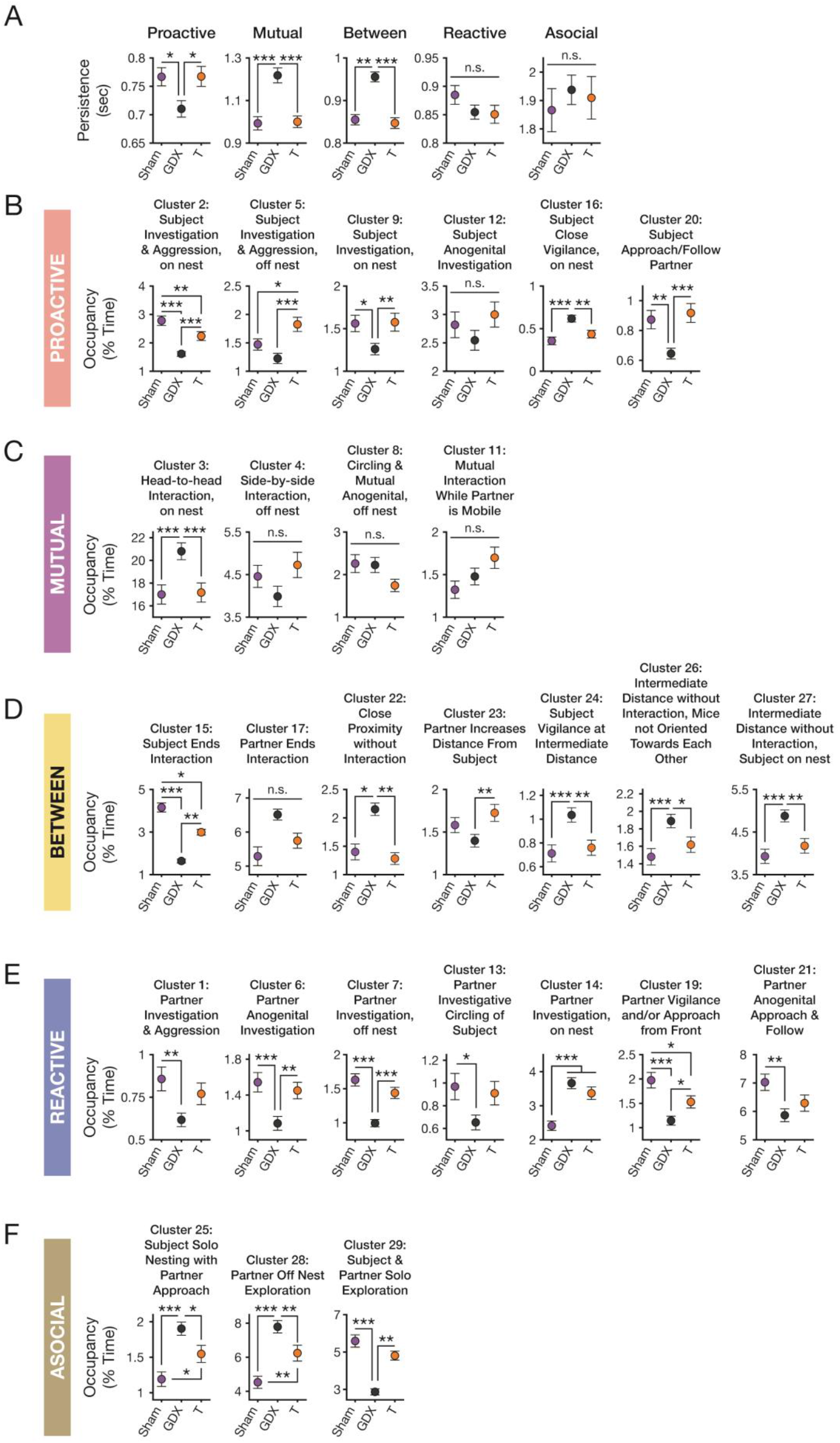
GDX disrupts occupancy and persistence of male behavioral action. A. Supercluster persistence as a function of hormonal manipulation. Proactive α_FDR_ = 0.0333; Mutual α_FDR_ = 0.0333; Between α_FDR_ = 0.0333; Reactive α_FDR_ = 0.05; Asocial α_FDR_ = 0.05. B. Occupancy across proactive clusters as a function of hormonal manipulation. Cluster 2 α_FDR_: 0.05; Cluster 5 α_FDR_: 0.0333; Cluster 9 α_FDR_: 0.0333; Cluster 12 α_FDR_: 0.05; Cluster 16 α_FDR_: 0.0333; Cluster 20 α_FDR_: 0.0333. C. Occupancy across mutual clusters as a function of hormonal manipulation. Cluster 3 α_FDR_: 0.0333; Cluster 4 α_FDR_: 0.0167; Cluster 8 α_FDR_: 0.05; Cluster 11 α_FDR_: 0.0167. D. Occupancy across between clusters as a function of hormonal manipulation. Cluster 15 α_FDR_: 0.05; Cluster 17 α_FDR_: 0.0167; Cluster 22 α_FDR_: 0.0333; Cluster 23 α_FDR_: 0.0167; Cluster 24 α_FDR_: 0.0333; Cluster 26 α_FDR_: 0.0333; Cluster 27 α_FDR_: 0.0333. E. Occupancy across reactive clusters as a function of hormonal manipulation. Cluster 1 α_FDR_: 0.0167; Cluster 6 α_FDR_: 0.0333; Cluster 7 α_FDR_: 0.0333; Cluster 13 α_FDR_: 0.0167; Cluster 14 α_FDR_: 0.0333; Cluster 19 α_FDR_: 0.05; Cluster 21 α_FDR_: 0.0167. F. Occupancy across asocial clusters as a function of hormonal manipulation. Cluster 25 α_FDR_: 0.05; Cluster 28 α_FDR_: 0.05; Cluster 29 α_FDR_: 0.0333. N = 144 sham sessions, 8 mice; N = 252 GDX sessions, 14 mice; N = 162 T sessions, 9 mice. Data presented as mean ± S.E.M. Linear mixed effects models were used for all statistical tests. Linear Mixed Effects Model formulas can be found in Methods. Cluster 10: Mutual anogenital behavior on the nest (Mutual Supercluster), Cluster 18: Initiate Anogenital (Proactive Supercluster), Cluster 30: Off nest exploration (Asocial Supercluster), and Cluster 31: Solo nesting (Asocial Supercluster) can be found in Fig. 5D and are omitted from this supplement. *: *p* < α_FDR_, **: *p* < 0.01, ***: *p* < 0.001.

## Methods

### Mice

All animal procedures were approved by the Princeton University Institutional Animal Care and Use Committee and were in accordance with National Institutes of Health Standards. Adult mice aged 8-22 weeks at the start of experiments were used for all studies. Mice were group-housed across all experiments with 2-5 same-sex mice per cage at 21-26 °C and 30-70% humidity in a 12-hour reversed dark-light cycle (OFF: 10:00 AM; ON: 10:00 PM). Experiments occurred exclusively during the subjective active (dark) phase. Food and water were provided *ad libitum*.

A total of 58 (N = 27 females, 31 males) subject mice were used in this study. For simultaneous behavior and multisite fiber photometry experiments, we used *Esr1*::*cre* (Jackson Laboratories stock number 017911, Maine, USA) male and female mice as experimental subjects, backcrossed to the CD1 strain (Charles River Strain Code 022, Massachusetts, USA). For behavior experiments without photometry, we used wildtype CD1 strain mice as subjects. Male and female subjects were used for all experiments. Sex was assigned by anogenital distance at weaning and confirmed visually via presence of testes (male) or ovaries (female) at gonadectomy or sham gonadectomy. All experimental subjects were sexually naive at the start of experiments. For mice used as social targets (partners), sexually naive wildtype CD1 strain males, CD1 strain females, and BALB/c strain males (Taconic Biosciences model number BALB-M, New York, USA) were used. All subject mice were naive to partner mice prior to interaction. All animals were bred within the Princeton University animal facilities or purchased directly from Charles River Laboratories or Taconic Biosciences.

The estrous cycle was monitored on experimental days for all female subjects and partner mice. Disposable pipettes filled with a small volume of distilled water were placed at the vaginal opening, and vaginal epithelial cells were collected by flushing the vagina with distilled water without inserting the pipette to avoid pseudopregnancy. Vaginal cytology was subsequently imaged under a brightfield microscope, and mice were staged based on the observed cell types. Mice with nucleated cells were assigned to proestrus, mice with cornified cells were assigned to estrus, and mice with leukocytes were assigned to diestrus. For comparisons between estrus and diestrus, data from proestrus and estrus days were combined.

### Custom Multisite Fiber Photometry

#### Implant

To record from 11 brain regions simultaneously, we developed a customizable multisite fiber photometry implant. Implants were designed in Autodesk Fusion360 (California, USA) to approximate dimensions of 11.5 mm X 9.5 mm X 5 mm (*length, width, depth*) with curved edges and skull facing surfaces to better match the contours of a mouse skull. Each implant was designed with 11 individual 200 μm through holes at defined stereotactic coordinates for fiber placement (see surgery coordinates below), 4 individual 1.2 mm through holes at cardinal positions in the design, and 4 insets for nuts in the corners of the implant. Implants were 3D printed by Protolabs (Minnesota, USA) using their MicroFine ABS-like plastic material.

To ensure through holes would fit fibers and were orthogonal to the implant surface, a custom holder was designed to hold the prints in a stereotaxic frame. The print was placed in this holder, leveled in the stereotaxic frame, and a 250 μm drill bit was used to clear any remaining 3D print material from the through holes. Next, 4 M0.6 brass hex nuts (MetricScrews.us, Mississippi, USA) were superglued into the printed insets. 200 μm optic fibers (FT200UMT, Thorlabs, New Jersey, USA) were cleaved to specified lengths (XL411, Thorlabs, New Jersey, USA), inserted into their matching through holes, and superglued in place. These lengths took into account final fiber targeting (see below) and the thickness of the implant. ArcN fibers were cut to a final length of 12 mm, and fibers for all other sites were cut to a final length relative to the ArcN fiber to match their final fiber depth (see below).

A custom polishing puck was built to fit the size of the 3D printed implants, and the top surface was polished using diamond lapping sheets (LF6D, LF3D, LF1D, LFCF2, Thorlabs, New Jersey, USA). Following polishing, implants were tested for efficiency using custom patchcords (see below) and only implants with efficiency > 70% were kept for experiments. Following efficiency testing, 4 alloy steel dowel pins (1.19 mm x 12.7 mm, McMaster-Carr 98381A983, Illinois, USA) were superglued into the pre-printed through holes, such that they were flush with the skull surface side of the implant.

#### Patchcord

To collect fluorescent emission from the photometry implant, we designed a custom multifiber branching-patchcord. This patchcord consists of two parts. First, we designed a custom 3D printed connector with 200 μm through holes at defined stereotactic coordinates to match the implants. This connector has approximate dimensions of 11.5 mm X 9.5 mm X 9.3 mm, 4 through holes to align implant and patchcord with the implants’ dowel pins, and 4 through holes at the corners of the connector for screw placement (Drive screws, Open Ephys, Lisbon, Portugal) to ensure a stable connection with the implant. Second, a custom branching-patchcord was purchased from Doric Lenses with 200 μm, 0.37 NA optic fiber (200/220/3000-0.37_5m_SMA-12xCL_LAF). The fibers were inserted into the through holes in the connector and superglued in place. After gluing, the custom polishing puck was used to polish the implant facing side of the patchcord.

### Surgery

For all procedures, mice were anesthetized (isoflurane: 3-5% induction; 1-2% for maintenance) and affixed to a stereotaxic frame prior to the initiation of surgery. Mouse skulls were leveled using confluence of the sagittal sinus and rostral rhinal vein (lateral vein that sits on the dorsal surface of the mouse brain between the olfactory bulb and prefrontal cortices) as the anterior position and lambda as the posterior position.

### Multisite Fiber Photometry

For two-color jRCaMP1b and GCaMP6f multisite fiber photometry recordings, 11 craniotomies were made to target SBN regions. At each of these craniotomies, *Esr1*::*cre* mice were injected with 300nl/site with a 1:1 mixture of pAAV1/Syn-Flex-NES-jRCaMP1b-WPRE-SV40 (titer: 2.7 x 10^13^; Addgene, Massachusetts, USA) and AAV2/EF1a-Cre-off-GCaMP6f-WPRE-hGHpA (titer: 1.9 x 10^14^) to drive expression of jRCaMP1b in ERα+ and GCaMP6f in ERα− neural populations, respectively.

Viral injection coordinates are as follows (in mm, relative to the confluence of the sagittal sinus and rostral rhinal vein; dorsoventral [DV] distance measured from brain surface): TTd, Anterior-posterior (AP): −1.40, Medial-lateral (ML): +0.35, DV: −3.10; LSv, AP: −2.60, ML: +0.60, DV: −3.35; MPOA, AP: −3.00, ML: −0.40, DV: −5.00; BNST, AP: −3.30, ML: +0.85, DV: −3.90; AHA, AP: −3.70, ML: −0.50, DV: −5.10; VMHvl, AP: −4.65, ML: +0.72, DV: −5.80; ArcN, AP: −4.70, ML: −0.33, DV: −5.90; MeA, AP: −4.70, ML: −2.25, DV: −5.20; PMv, AP: −5.50, ML: −0.58, DV: −5.5.; PA, AP: −5.50, ML: +2.40, DV: −5.00; lPAG, AP: −7.80, ML: +0.50, DV: −1.90.

Subsequently, the custom multisite photometry implant was lowered with all fibers inserted into their matching craniotomies to a depth of −5.55 mm. This produced effective fiber depths (in mm relative to brain surface) as follows: TTd: −2.85; LSv: −2.90; MPOA: −4.55; BNST: −3.50; AHA: −4.60; VMHvl: −5.50; ArcN: −5.55; MeA: −4.80; PMv: −5.4; PA: −1.80; lPAG: −1.60.

Implants were fixed to the skull with Metabond (Parkell, New York, USA). Animals were allowed to recover for a minimum of 2 weeks.

### Gonadectomy

To reduce circulating gonadal steroid hormones, orchidectomies were performed in mice with testes and ovariectomies were performed in mice with ovaries. For orchidectomies, an abdominal approach was taken. A single incision was made in the skin and another through the abdominal muscles. The testes were then removed from the internal body cavity and their attached blood vessels ligated with absorbable suture (Coated Vicryl, Ethicon, Johnson & Johnson, New Jersey, USA). A cut was then made to the testis side of the ligation to remove each testis. The internal muscles were then sutured using absorbable sutures, and the skin incision was closed with wound staples. For ovariectomies, a lateral flank approach was taken. Single incisions were made along the left and right flanks of the mouse and the underlying muscle wall. The ovaries were then removed from the body cavity, and the connection between the ovary and its corresponding uterine horn was ligated with absorbable suture. A cut was then made to remove each ovary. Wounds were closed as in orchidectomy.

For all sham surgeries, incisions were made as in real orchidectomies and ovariectomies, testes or ovaries were visually identified, and then wounds were closed as described above. Wound staples were removed following wound healing and within 10 days of surgery. All mice recovered for two weeks prior to the initiation of post-gonadectomy experiments.

### Osmotic minipump implantation

During the same surgery as gonadectomy, an osmotic minipump (Alzet Osmotic Pump Model #1004, California, USA) was implanted subcutaneously between the scapulae. Briefly, an incision was made between the shoulders, and the pump was inserted into the interstitial space between the skin and muscle. Then the wound was sutured using absorbable suture. The mini-pump was filled with either a sesame oil vehicle or 60 μg testosterone in sesame oil for delivery of 6.6 μg/hr testosterone.

### Behavioral assays

All animals underwent the same social interaction assays. Briefly, subject mice were allowed to explore a cage for 1 minute before introduction of a partner mouse for a 4-minute interaction. For behavior only experiments, subject mice were placed into the cage at the start of the recording. For multisite fiber photometry experiments, mice were placed on a platform outside the cage at the start of the recording and allowed to enter the cage on their own. Each day, subject mice underwent 6 interactions across all possible combinations of partner (CD1 male, CD1 female, and Balb/c male) and territory (homecage, partner’s cage) conditions in a random order.

Subject mice interact with the same partner mouse in their homecage and their partner’s cage. Nests and food pellets were left in the cage during the interaction. Black paper-chip bedding was used during housing and experiments to enhance contrast for behavior imaging. Experiments were replicated on 3 non-sequential days before and after gonadectomy or sham surgery. Each day, subject mice interacted with novel partners.

### Data acquisition

All code was custom written MATLAB code unless otherwise specified. In total, data acquisition and analysis used custom code written in Bonsai (Bonsai Foundation, Wales and England, UK), MATLAB (Mathworks, Massachusetts, USA), and Python (Python Software Foundation, Delaware, USA).

### Behavioral video acquisition

All behavior was recorded using a top and side camera (40Hz, FLIR, SpinView). The top camera was centered 50 cm above the cage, and the side camera was positioned at a height of 30 cm. Using MATLAB’s Stereo Camera Calibrator application and a charuco board, we corrected camera images for fisheye distortion of images and to convert pixels into cm coordinates.

### Fiber photometry data acquisition

Multisite fiber photometry recordings were made using an FP3002 (Neurophotometrics, California, USA). The FP3002 contains 415, 470, and 560 nm LEDs for isosbestic, GCaMP and RCaMP excitation, respectively. A dichroic mirror splits GCaMP and RCaMP emission into separate channels where they are further filtered with bandpass filters (green channel: 494-531 nm; red channel: 586-627 nm). Mirrors reflect this emission onto distinct locations on the sensor of a CMOS camera. Video was acquired at 120 Hz with 415, 470, and 560 nm LED stimulation interleaved across all frames. Custom Bonsai code controlled the FP3002 system, and the Bonsai software was driven by an external data acquisition device.

### Data synchronization

Data acquisition was controlled via a MC USB-200 series DAQ with DAQami software (Measurement Computing, Massachusetts, USA). A TTL pulse was sent via the DAQ to start and stop behavioral video and fiber photometry recordings. DAQami recorded timestamps for FP3002 start and stop, as well as each frame capture for behavioral cameras. Bonsai software for fiber photometry data acquisition recorded timestamps for each frame capture for the FP3002 system. These timestamps were used to align fiber photometry and behavioral video data.

### Behavioral analysis

#### Pose tracking

We used SLEAP (v1.3.1) to track mouse pose and nest location during social interactions. We built our mouse model using 6551 frames from 163 different social interactions. We built our nest model using 1904 frames from 89 different social interactions. Training was conducted using 80:20 train:test splits. For subject and partner mice, we tracked the following points: nose, head, neck, left and right ear, left and right forepaws, trunk, left and right hindpaws, and tailbase. Mouse centroid was defined as the mean of the *xy* coordinates of all tracked nodes. For nests, we tracked the center of the nest. Mice were tracked using a top-down model, and nests were tracked with a bottom-up model.

#### Feature definition and preprocessing

Prior to any computation of behavior features, lens correction was performed on behavior video frames using the MATLAB function ‘undisortPoints’ and camera calibration statistics derived from MATLAB’s Stereo Camera Calibrator. Tracked mouse posture and nest location were used to generate 78 unique behavior features.

These features can be found in Table S1. Individual mouse egocentric coordinates were normalized by body length on a per-session basis. Tracked pose coordinates and all behavior features were preprocessed in the following manner, on a session-by-session level: (1) outliers were detected at values ≥±3 standard deviations from the mean coordinate or feature value and were replaced with a value of ‘NaN’; (2) ‘NaN’ values were replaced via piecewise cubic interpolation; (3) interpolated signal was smoothed using a 10^th^ order median filter followed by a Gaussian filter of 0.25 seconds.

### Unsupervised behavior analysis

#### Behavior map and cluster generation

To comprehensively capture the behavioral landscape during mouse social interaction, we adapted previous unsupervised behavior classification strategies^32,33^ that define behaviors as high density clusters in a two dimensional embedding of behavior features. To achieve dense clusters, we embedded behavior features from the first 50 experimental subject mice (1,800 social interaction sessions, 18.1 M frames) into two dimensions using tSNE. To ensure all 78 behavior features existed within a common coordinate space, features from these 50 mice were *z*-scored prior to any dimensionality reduction. Mean and standard deviation from all features, as well as PCA loadings for egocentric coordinates, were saved to allow subsequently collected data to be added to the embedding. To embed our data in two dimensional tSNE space, we took an approach known as importance sampling that allowed us to capture more rarely expressed behaviors. This process involves two rounds of tSNE embedding.

First, using custom python code, approximately 100,000 frames were uniformly sampled across the 1,800 videos, and their behavior feature data were embedded into a two-dimensional tSNE (sklearn.manifold.TSNE, perplexity = 100, early_exaggeration = 12, init = ‘pca’, metric = ‘cosine’). To map the remaining ∼18 M frames from 78-dimensional behavior feature space to two dimensional tSNE space, a separate multilayer perceptron regression model (MLPR) was built (sklearn.neural_network.MLPRegressor, hidden layer size of 800 x 700 x 600 x 500 x 400 units, L2 regularization alpha set to 0.001). Next, all 18.1 M frames were binned into a 128 x 128 probability density histogram with a two-dimensional gaussian kernel of 3 standard deviations. We then used watershed clustering (skimage.segmentation.watershed) to identify 35 unique clusters from the smoothed probability density histogram.

Subsequently, to generate the final tSNE embedding, 2 random frames from each of the 35 clusters were sampled from each of the 1800 videos. Samples were not taken from videos that lacked frames from the given cluster. This resulted in an importance sampled dataset of approximately 125,000 frames across 1,800 sessions, or approximately 3,600 frames per cluster. Next, we embedded the behavior features from these importance sampled frames into two dimensional tSNE space as described above and used the MLPR to embed the remaining 18 M frames.

To create a final two-dimensional probability density histogram, or “behavior map”, we ran the MATLAB function ‘histogram2’ on the x-y coordinates of all 18.1 M frames of tSNE embedded behavior video data. The number of bins was determined using the Freedman-Diaconis rule, which is less sensitive to outliers, and was set at 184 x 184. This histogram was then smoothed using a gaussian kernel of 2.5 standard deviation. Pixels with probability density < 1 x 10^−6^ were set to 0, low prominence peaks (probability difference < 1 x 10^−5^) were suppressed using the function ‘imhmax’, and the behavior map was renormalized such that the sum of its pixels equaled 1. We used the function watershed to generate cluster boundaries based on local density maxima, yielding 31 putative behavior clusters. We set a 150 ms minimum cluster duration. Frames that persisted in a cluster for less than or equal to 150 ms were reassigned via nearest neighbor interpolation using the MATLAB function ‘interp1’.

Behavior features collected from mice after the 50 used to generate the behavior map were converted to the common *z*-score space using the mean and standard deviations of the behavior features from the first 50 mice. Mouse egocentric posture was converted to PCA space using the PCA loadings from the first 50 mice and then converted to *z*-score as described above. We then used the MLPR described above to embed these data into the common tSNE behavior space. In total, 20,845,987 frames from 2,000 social interaction sessions across 58 mice were embedded into the tSNE behavior space.

Cluster occupancy was defined, on a per-session basis, as the percentage of frames belonging to a given cluster out of the total number of frames for the behavioral interaction session. Cluster persistence was defined as the duration of a single cluster bout.

#### Supercluster generation

To identify classes of similar behavioral actions, we generated “superclusters” of the tSNE derived clusters. To achieve this, we first generated the mean value of all behavior features, except for egocentric posture PCs, on a per cluster basis. Then, we hierarchically clustered the clusters based on their *z-*scored mean feature values using Ward’s method. This yielded 5 superclusters, consisting of clusters with similar behavior features.

Supercluster occupancy was defined, on a per-session basis, as the percentage of frames belonging to a given supercluster out of the total number of frames for the behavioral interaction session. Supercluster persistence was defined as the duration of a single supercluster bout.

### Comparison of unsupervised behavior maps

To determine if behavior occupancy is predictive of sex, social context, or hormone state, we used binomial or multinomial-softmax regression and a leave-one-session-out cross-validated (LOOCV) approach to classify session identity. For each session, we generated vectors of cluster (31 x 1) and supercluster (5 x 1) occupancy for each session. We built separate cluster and supercluster classifiers. To build these classifiers, we built cluster and supercluster training matrices of size N-sessions x 31 or 5. Since occupancy across clusters or superclusters sums to 1 for each session, to avoid issues of collinearity, we used PCA to transform these matrices. We included all PCs up to 99% explained variance and *z*-scored each PC, producing a final training matrix of size N-sessions x N-PCs. Using this matrix and a vector of real labels, we trained the LOOCV classifiers. For LOOCV, the model is trained iteratively using data from all sessions but one and predicts the single held-out session. We used the full set of predictions to generate confusion matrices and compute F1 score, precision, and recall for the models. Separately, we generated a chance distribution to determine statistical significance. Briefly, instead of training using real session labels, session identity labels were shuffled 1,000 times, and a separate LOOCV model built for each shuffled vector. F1 score was computed for each model trained on shuffled labels.

In females, to determine if the relationship between social context and behavior differed across estrous, we trained a 10-fold CV classifier on either real or shuffled cluster or supercluster occupancy from sessions when the mouse was in diestrus. For each fold of training, in addition to testing on the diestrus test set, we predicted territory or partner identity from the sessions when the mouse was in estrus using the real and shuffled models. F1 score was computed for real and shuffled models.

In males, to determine if the relationship between social context and behavior changed following GDX, we trained a 10-fold CV classifier on supercluster occupancy from pre-GDX sessions. For each fold of training, in addition to testing on the held-out pre-GDX test set, we predicted territory or partner identity from post-hormonal manipulation behavior sessions from the sham, GDX, and T groups. F1 score was computed for all model results.

To determine if GDX changed behavior maps more than sham GDX or GDX + T, we used Jensen-Shannon (JS) divergence to compare cluster occupancy between behavior maps. JS divergence measures the similarity between two probability distributions (*i*.*e*., cluster occupancy) with more similar distributions approaching a JS divergence value of 0. Given two probability distributions *P* and *Q*, JS divergence is defined as,

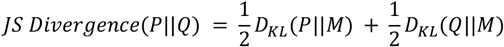

where,

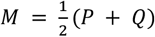

and D_KL_ is the Kullback-Leibler divergence defined as,

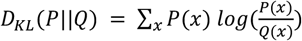

To quantify change in behavior map from pre-to-post hormonal manipulation, the cluster distributions of the three pre-GDX behavior sessions of a given social context were compared to the three post-GDX behavior sessions to generate 9 comparisons per mouse. To quantify the change in behavior map across estrous, cluster distributions from estrus and proestrus behavior sessions were compared to diestrus behavior sessions from the same mouse. Only data from pre-GDX female mice was used to quantify JS divergence across estrous.

### Calcium imaging analysis

#### Data processing

Calcium imaging data was processed using custom MATLAB (Massachusetts, USA) and Python (Delaware, USA) code. First, data from both green and red channels were de-interleaved to separate the emitted signal in response to each of the three LEDs, and the timestamps of each signal were aligned to behavioral video timestamps. Next, GCaMP, RCaMP, and isosbestic signals were linearly detrended and regressed to baseline by fitting and subtracting a single exponential decay from them. ΔF/F was computed as 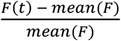. Subsequently, TMAC^38^ was used to remove motion artifacts and gaussian noise. For each fiber, GCaMP and RCaMP signals were corrected using both the red and green channel emission in response to 415 nm stimulation of that fiber. Whichever of the two corrected signals (i.e., red or green channel isosbestic corrected) had a lower noise level, defined as its standard deviation, was chosen as final TMAC-derived signal for that recording. Following TMAC, data were filtered using a 3^rd^ order Savitsky-Golay filter with a one second window, and then quality control was performed on the data. Signal was excluded at a single session by fiber basis based on the following criteria: 1) no detectable calcium transients of at least 1% ΔF/F (using MATLAB function ‘findpeaks’); 2) high correlation between corrected signal and isosbestic (r^2^ > 0.8); 3) mechanical signal distortion due to torque on patchcord. 97.7% of 15,026 signals passed quality control. Finally, GCaMP and RCaMP signals from each fiber were concatenated and z-scored on a per day basis. For analyses that relied on all signals from an animal on a given session, excluded data were replaced with z-scored gaussian noise with standard deviation set to the standard deviation from that fiber for that day’s sessions.

#### Projecting neural signals onto behavior maps

To project neural activity onto behavior maps to generate neural activity maps, we indexed each frame of neural activity data to its corresponding x-y position in tSNE space. Then, on a per fiber, per session basis the mean *z*-scored ΔF/F neural population activity was computed for each x-y position in tSNE space. Finally, these maps were smoothed using a 2D gaussian kernel of 2.5 standard deviations.

#### Quantification of mean population activity during clusters and superclusters

For comparing mean *z*-scored ΔF/F during a given cluster or supercluster, only neural signals that passed quality control described above were used for analysis. For each neural population that passed quality control, all frames belonging to a given cluster or supercluster were identified and neural population activity was averaged on a per session basis.

#### Social context and action decoders

To decode social context and action, we built 10-fold CV classifiers using binomial (territory) or multinomial (partner, social action) regression. Briefly, for each mouse, a balanced input matrix of neural population activity by time was generated by randomly sampling an equal number of imaging frames from either each territory, partner, or supercluster, and a corresponding label vector of territory, partner, or supercluster labels was generated. The input matrix and corresponding label vector were built using data from across all pre-GDX days for each mouse for all decoders, but this was done separately for each conjunctive territory-partner social context for the social action decoder

To determine model performance relative to chance, separate classifiers were trained using both real and shuffled label vectors.

To determine the contribution of ERα+ and ERα− neural populations to social context or action encoding, separate 10-fold CV models were trained using an input matrix where all ERα+ and ERα− populations were shuffled iteratively. F1 loss was defined as the difference between the real model and the shuffled model F1 scores.

To determine whether the social context or action code was modular or distributed, we rank-sorted the model weights across neural populations from the original model trained on real labels. For multinomial models, weights were averaged across comparison axes prior to rank sorting. Then, we iteratively trained 10-fold CV models using an input matrix where neural populations were cumulatively shuffled in order of descending weight with each iteration, starting with the population with the greatest predictive power in the original model.

The input matrix for the supercluster decoder was split into 11 matrices of equal duration. 10 of these segments were used to train the 10-fold CV models discussed above. To determine the context-specificity of the social action code, during each fold of training, the model was tested on both the test set for that fold as well as the held-out 11th segment from each of the 6 conjunctive territory-partner social contexts.

#### Territorial regression analysis

To build a linear model explaining the relationship between social context and behavior-associated neural activity, we first generated vectors of mean *z*-scored neural population activity during each cluster (1 x Clusters) within each day for each neural population. As not all clusters were sampled during each behavior session, these vectors were averaged across days to generate a single vector of neural activity during each behavior cluster for each neural population from each mouse in each of the 6 conjunctive territory-partner contexts. To perform analyses on the code across all partners, these vectors were concatenated across the 3 partner contexts within either the home or partner’s territory. To determine the requirement of proactive behaviors in the observed rescaling, these vectors were generated both with and without proactive clusters included.

Next, using only signals that passed quality control, we regressed the home territory vector to the partner territory vector using the MATLAB function ‘regress’. We did this for each of the 3 social partner contexts as well as for the full home and partner’s territory vectors concatenated across all partners. Vectors were regressed to the corresponding vector of the same neural population recorded from the same mouse. Territory regression analysis was performed separately for animals pre-GDX and post-GDX/Sham/T. To determine if the observed regression results differed from chance, we computed 10 shuffled regressions for each pair of home and partner’s territory vectors.

#### Identification of neural subnetworks

To identify subnetworks of neural populations within the SBN based on the relationship between the neural action code across territories, we hierarchically clustered matrices of territory regression model coefficients and intercepts (Neural Populations X Partners*2). To identify subnetworks of neural populations within the SBN based on GDX-induced, population-specific changes to the relationship between the neural action code across territories, we first generated mean matrices of territory model coefficients and intercepts for data from sham or GDX mice. To identify the subnetworks, we hierarchically clustered the GDX - Sham difference matrix (Neural Populations X Partners*2).

#### Cosine distance comparison of neural signals

To test how GDX impacted the relationship of the SBN neural action code between partners, we used cosine distance to quantify the similarity of neural representations of behavior across contexts. Specifically, we used the vectors of mean *z*-scored neural population activity (1 x Clusters) generated during the territorial regression analysis to compare AGG+ to AGG- or FEM sessions. Cosine distance was computed using the equation:

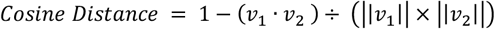

Where *v*_1_ corresponds to the AGG+ vector, *v*_2_ to the AGG- or FEM vector, and ‖‖*v*_*n*_‖‖ to the vector magnitude, or euclidean norm. We compared vectors from both within and across mice within the same hormonal manipulation (*i*.*e*., sham or GDX) to compute cosine distance between vectors from AGG+ and AGG- or FEM contexts. All cosine distance comparisons were done within a territory context (*i*.*e*., home or partner’s territory).

#### Histology for virus expression and fiber placement

Following the completion of experiments, animals were deeply anesthetized with euthaphen (sodium pentabarbitol, Dechra, England, UK) and perfused with a 4% paraformaldehyde (PFA) dissolved in 1x Phosphate Buffer Solution (PBS). Animals were decapitated and skulls with implants intact were allowed to post-fix in 4% PFA in 1x PBS for at least 3 days to ensure visible fiber tracts following cyrosectioning. After post-fixing with the skull intact, the brain was removed and post-fixed for additional 12-24 hours in 4% PFA in 1x PBS. After this final post-fix, brains were moved to 30% sucrose in 1x PBS for cryoprotection. Once brains became denser than the sucrose solution, they were embedded in optimal cutting temperature mounting medium (Fisher Healthcare Tissue-Plus OCT, Fisher Scientific, New Hampshire, USA) and frozen over dry ice. 50 μm cryosections of frozen tissue were collected and stored in 1x PBS prior to mounting. Slices were then washed in PBS and mounted on slides, allowed to dry, coverslipped with a mounting medium (EMS Immuno Mount DAPI and DABSCO, Electron Microscopy Sciences, Pennsylvania, USA), and clear nail polish was used to seal the coverslip to the slide. After at least 12 hours of drive, slides were imaged with a digital robotic slide scanner (NanoZoomer S60, C13210-01, Hamamatsu Photonics, Hamamatsu, Japan) and figures were generated using QuPath^71^ and ImageJ (NIH, Washington, D.C., USA). Fiber tip locations were manually identified and localized using the Allen Mouse P56 Brain Atlas (atlas.brain-map.org, Washington, USA).

### Inferential statistical analyses

For all statistical tests, *p* values were compared against an α significance threshold of 0.05. Where necessary, we corrected for multiple comparisons using the false discovery rate (FDR) method^72^ to adjust the alpha threshold for significance. For linear mixed effects models (LMEs) described below, a unique LME was generated for each possible comparator, and *p* values were saved for each unique comparison. These *p* values were then used to determine the FDR-corrected significance threshold. LMEs were built using the MATLAB function ‘fitlme’ with ‘FitMethod’ set to ‘REML’.

### Comparison of unsupervised behavioral landscapes

To determine if LOOCV models could significantly decode relative to chance, we computed *p*-value for each model’s performance as the proportion of shuffled model F1 scores greater than or equal to the real model’s F1 score.

To determine if models trained on diestrus behavioral occupancy could decode social context on diestrus or estrus days, we used a paired t-test to compare F1 scores between real and shuffled model results.

To determine if models trained on male pre-GDX behavioral occupancy performed better decoding social context from the behavioral occupancy of Sham, GDX, or T mice, we built an LME with the following formula: ‘F1 ∼ HormoneState + (1|CV)’.

To compare JSD across multiple hormonal manipulations, we built an LME with the following formula: ‘JSD ∼ HormoneState + (1|Mouse)’.

### Comparison of cluster or supercluster occupancy and persistence

To test if social context determined cluster or supercluster occupancy, we built separate LMEs with the following formulas: ‘Occupancy ∼ Territory * Partner + Day + (1|Mouse)’ or ‘Occupancy ∼ Territory * Partner + (1|Mouse)’. After we found no effect of ‘Day’ on the occupancy of superclusters in any context, we used the second formula for all tests.

To test if cluster or supercluster occupancy differed across sex, we built separate LMEs with the following formulas: ‘Occupancy ∼ Sex + (Partner|Territory) + (1|Mouse).’ To test if cluster or supercluster occupancy differed across territories, we built separate LMEs with the following formulas: ‘Occupancy ∼ Territory + (1|Partner) + (1|Mouse).’ To test if cluster or supercluster occupancy differed across partners, we built separate LMEs with the following formulas: ‘Occupancy ∼ Partner + (1|Territory) + (1|Mouse).’ To test if proactive occupancy differed across estrous stage, we built an LME with the following formula: ‘Occupancy ∼ Estrous + (Partner|Territory) + (1|Mouse).’ To test if hormone state determined cluster or supercluster occupancy or persistence, we built separate LMEs with the following formulas: ‘Occupancy ∼ HormoneState + (1|Mouse)’ or ‘Persistence ∼ HormoneState + (1|Mouse)’.

### Distance to nest center across hormone states

To determine if hormonal state was predictive of distance to nest, we computed the mean distance to nest centroid for all male behavioral sessions post-manipulation. We then compared these distances separately within the home and partner’s territory using separate LMEs with the formula: ‘Distance ∼ HormoneState + (1|Mouse)’.

### Comparison of supercluster neural activity levels

To test if social context determined neural activity during different superclusters, we built separate LMEs with the following formula: ‘Zscore ∼ Territory * Partner + (1|Population) + (1|Mouse).’ To test if behavior-associated neural activity levels changed following GDX, we built LMEs to compare *z*-scored neural activity within each supercluster in GDX and sham males using the formula: ‘Zscore ∼ HormoneState + (1|Population)’.

To determine whether neural activity in the home or partner territory was predictive of the pattern of behavioral occupancy across partners, we built separate LMEs for each supercluster-territory conjunctive pairing with the formula: ‘Occupancy ∼ Zscore + (1|Population) + (1|Mouse) + (1|Partner)’.

### Social context and action decoders

To test if models performed significantly differently than chance, F1 scores from real and shuffled models were compared using a paired t-test. We used a paired t-test to determine if F1 loss significantly differed when ERα+ or ERα− populations were shuffled. To determine when the cumulatively shuffled decoder reached chance levels, we built an LME comparing F1 score when all 22 populations were shuffled to the F1 score from the fully unshuffled model and all other cumulative iterations from 1-21 populations shuffled. We used the following formula for these LMEs: ‘F1 ∼ CumulativeIteration + (1|Mouse)’.

### Territorial regression analyses

To test if regression statistics between models (*i*.*e*., real vs shuffle, male vs female, sham ERα+ vs sham ERα− vs GDX ERα+ vs GDX ERα−), we compared distributions of real and shuffled model coefficients, intercepts, and R^2^ values using a two-sample Kolmogorov-Smirnov test. If more than two samples were compared, each possible comparison was made and the α threshold for significance was adjusted using the FDR method.

To determine if model residuals were significantly different from chance on a cluster-by-cluster basis, the model residuals for each cluster were compared to the shuffled model residuals using the following formula: ‘Residual ∼ Shuffle + (1|Population) + (1|Mouse)’. To determine if model coefficients or intercepts were significantly different from chance on a population-by-population level, real coefficients and intercepts were compared to shuffled model coefficients and intercepts using the following formulas: ‘Coefficient ∼ Shuffle + (1|Mouse)’ and ‘Intercept ∼ Shuffle + (1|Mouse)’.

### Comparison of cosine distance

To test if GDX impacted the similarity between neural representations of behavior during interactions between AGG+ and AGG- or FEM partners, we built separate LMEs with the following formula: ‘CosineDistance ∼ HormoneState + (1|Population).’

## Notes

### Competing Interest Statement

The authors have declared no competing interest.

